# Genuine Selective Caspase-2 Inhibition with new Irreversible Small Peptidomimetics

**DOI:** 10.1101/2021.12.13.472505

**Authors:** Elodie Bosc, Julie Anastasie, Feryel Soualmia, Pascale Coric, Ju Youn Kim, Gullen Lacin, Eric Duplus, Philippe Tixador, Bernard Brugg, Michelle Reboud-Ravaux, Serge Bouaziz, Michael Karin, Chahrazade El Amri, Etienne Jacotot

**Author notes:** **Correspondence should be addressed to:** Prof Etienne D. Jacotot, PhD, Department of Pathology & Cell Biology, Taub Institute for Research on Alzheimer’s Disease and the Aging Brain, Columbia University Medical Center, New York, New York 10033.

## Abstract

Caspase-2 (Casp2) is a promising therapeutic target in several human diseases including nonalcoholic steatohepatitis (NASH) and Alzheimer’s disease (AD). However, the design of active-site-directed inhibitor selective to individual caspase family members is challenging because caspases have extremely similar active sites. Here we present new peptidomimetics derived from the VDVAD pentapeptide structure, harboring non-natural modifications at the P2 position and an irreversible warhead. Enzyme kinetics shows that these new compounds, such as LJ2 or its specific isomer LJ2a, and LJ3a, strongly and irreversibly inhibit Casp2 with genuine selectivity. According to Casp2 role in cellular stress responses, LJ2 inhibits cell death induced by microtubule destabilization or hydroxamic acid-based deacetylase inhibition. The most potent peptidomimetic, LJ2a, inhibits human Casp2 with a remarkably high inactivation rate (k_3_/K_i_ ~ 5 500 000 M^−1^s^−1^) and the most selective inhibitor, LJ3a, has a near to 1000 times higher inactivation rate on Casp2 as compared to Casp3. Structural analysis of LJ3a shows that spatial configuration of C_α_ at the P2 position determines inhibitor efficacy. In transfected human cell lines overexpressing site-1 protease (S1P), sterol regulatory element-binding protein 2 (SREBP2) and Casp2, LJ2a and LJ3a fully inhibit SREBP2 activation, suggesting a potential to prevent NASH development. Furthermore, in primary hippocampal neurons treated with β-amyloid oligomers, submicromolar concentrations of LJ2a and of LJ3a prevent synapse loss, indicating a potential for further investigations in AD treatment.

## Introduction

Cysteine-dependent aspartate-specific proteases (Caspases) are a family of cysteine endoproteases (C14A family, CD clan), unique to the animal kingdom [1], and are well known as central effectors and regulators of apoptosis and inflammation [2]. They are also involved in the regulation of non-apoptotic cell death pathways, as well as in various physiological processes which include proliferation, differentiation, cell migration, several functions of the nervous system (e.g., synaptic plasticity, axonal guidance, long-term potentiation, pruning of dendritic spines), cell cycle control, and stress responses [3]. These functions are critical and indicate that Caspases are therapeutic targets of great interest for several diseases [4, 5, 6].

Caspase-2 (Casp2, Nedd2, Ich1), the most evolutionary conserved member of the Caspase family, has numerous interesting properties and functions, which suggest that its specific targeting could lead to high potential drug candidates [7, 8]. First, it is non-essential for physiological programmed cell death [9] whereas it mediates stress-induced and pathologically induced apoptosis [10]. Casp2 is involved in the response to a wide panel of stresses (endogenous, infectious, physicochemical, xenobiotic, metabolic, inflammatory) by initiating apoptotic cell death pathways, by repressing autophagy, or by activating the inflammasome [7–10]. At the hepatic level, Casp2 plays an essential role in the pathogenesis of non-alcoholic steatohepatitis (NASH) [11, 12]. Within the nervous system, it is involved in synaptic plasticity and cognitive flexibility [13], and several neuropathological mechanisms such as neonatal brain damage [14], retinal ischemia [15], the synaptotoxic effects of β-amyloid peptide [16, 17], and tauopathies [18]. Casp2 is required for the cognitive decline seen in human amyloid precursor protein transgenic mice and experiments with Casp2 deficient mice implicate Casp2 as key driver of synaptic dysfunction in AD [17]. Consequently, there is a need to design Casp2 selective inhibitors (or substrates) to precisely define its apoptotic and non-apoptotic roles and to interfere pharmacologically with disease-related pathways.

Caspases cleave peptides and proteins after an Asp (P1 position) [1]. This unique property among cysteine-proteases, has led to the development of hundreds of active site-directed small peptides and peptidomimetics that are Caspase-specific [19]. Indeed, medicinal chemistry of Caspase inhibitors led to potent druggable peptide derivatives (e.g., Q-VD-OPh, an irreversible pan-caspase inhibitor [20]), to the clinical development of potent peptidomimetics (e.g., Belnacasan/VX765 [21], a reversible inhibitor of inflammatory caspases), and to the advanced clinical development of safe broad-spectrum caspase inhibitor (e.g., Emricasan [22], an irreversible pan-caspase inhibitor) [23]. However, the design of active site-directed inhibitors selective of one given individual Caspase is highly challenging because caspases have similar active sites, and most Asp^P1^-containing small peptides are efficiently recognized by several caspases [24]. This is even more critical when designing Casp2 inhibitors, as Casp2 and Caspase-3 (Casp3) share highly similar features regarding their active sites and inhibition by synthetic substrates [25, 26]. Studies with synthetic peptides containing C-terminal reversible (aldehyde; CHO) or irreversible (fluoromethylketone; fmk) warheads have established that Casp2, Casp3, and Casp7 preferentially cleave substrate having the general structure X^P5^-Asp^P4^-X^P3^-X^P2^-Asp^P1^-CHO/fmk (with relative permissivity at P5, P3 and P2), whereas other Caspases poorly recognize Asp^P4^-containing peptides [27–29]. Casp2 requires the presence of a P5 residue to recognize peptide substrates whereas Casp3 and Casp7 do not [19, 29]. Accordingly, the well-defined pentapeptide-based inhibitors of Casp2 (i.e., Ac-VDVAD-CHO, z-VDVAD-fmk, Q-VDVAD-OPh) are also efficient inhibitors of Casp3 [30, 31].

Using structural information on the Casp2 and Casp3 active sites and molecular modeling, Maillard et al., identified that the replacement of Alanine (P2 position), in the non-selective Ac-VDVAD-CHO peptide, by bulky residues, for instance a substituted isoquinoline or a 3-(S)-substituted proline, resulted in peptides with 20 to 60-fold increased selectivity for Casp-2 [32].

We further elaborated from the Maillard and coworkers’ approach and designed a pentapeptide derivative, named LJ2, which combined a N-terminal quinolyl-carbonyl and a C-terminal difluorophenoxymethylketone warhead with the Casp2-preferred pentapeptide backbone VDVAD where the Ala in position P2 was replaced by a substituted isoquinoline [33]. Afterward, using a hybrid combinatorial peptide substrate library, Poreba et al reported a peptidomimetic series having the general structure X^P5^-Glu^P4^-X^P3^-Ser^P2^-Asp^P1^-OPh where X^P3^ is a substituted Thr (*O*-benzyl-*L*-allothreonine), and X^P5^ is a 2-carboxy-indoline [34]. However, these irreversible inhibitors showed significant but still moderate selectivity for Casp2, as the best compound of the series, NH-23-C2, showed K_obs_/I ratio vis-a-vis Casp3 being ~32 times lower than for Casp2) [34].

Here we present the structure and characterization of irreversible peptidomimetics having the general structure Quinadoyl-Val^P5^-Asp^P4^-Val^P3^-X^P2^-Asp^P1^-difluorophenoxy-methyl-ketone, with X^P2^ being either 6-methyl-tetrahydro-isoquinoline (LJ2, LJ2a, and LJ2b) or a 3-(S)-neopentyl proline (LJ3a). Kinetics using recombinant human Casp2 and Casp3 show that, these inhibitors have very strong off-rate (k3/Ki) and selectivity toward Casp2. Particularly, LJ3a is highly selective for Casp2 (946 times less efficient on Casp3), far above previously described Casp2 inhibitors [32, 34]. We further show that these potent and selective Casp2 inhibitors have strong effects in several biological models including protection against cell death induced by microtubule destabilization, blockage of Casp2- and S1P-mediated SREBP2 activation and inhibition of synapse loss in primary neurons treated with β-amyloid oligomers.

## Materials and methods

### Enzymes, substrates, and inhibitors

The human active recombinants caspase-2 (#ALX-201-057), caspase-6 (#BML-SE170), and caspase-1 (#ALX-201-056) were purchased from Enzo Life sciences, Inc (New York, USA). The human active recombinant Caspase-3 (#707-C3) was purchased from R&D Systems (Minneapolis, USA). Purified Human active Cathepsin-B (#BML-SE198), Cathepsin-D (#BML-SE199), and Cathepsin-L (#BML-SE201) were purchased from Enzo Life sciences. The human active Plasmin, Thrombin, and Trypsin were purchased from Sigma-Aldrich. Human Kallikrein-1 (#2337-SE-010), Kallikrein-6 (#5164-SE-010), Kallikrein-8/Neuropsin Protein (#2025-SE-010) were purchased from R&D systems. Stock solutions were stored at −80°C. The fluorogenic substrates Ac-VDVAD-AMC, Ac-DEVD-AMC, Ac-VEID-AMC, z-RR-AMC, RLR-AMC and the FRET substrate Phe-Arg-Leu-Lys(Dnp)-D-Arg-NH2, as well as the reversible Inhibitors, Ac-DEVD-CHO (#ALX-260-030) and Ac-VDVAD-CHO (#ALX-260-058) were purchased from Enzo Life sciences, Inc (New York, USA). The fluorogenic substrates Boc-QAR-AMC and Boc-VPR-AMC were purchased from Bachem AG (Bubendorf, Switzerland). The compounds c33, h33, k33, and q33 were obtained from Dr Michel Maillard, CHDI Foundation, Inc. (New York, NY). Broad spectrum inhibitors, z-VAD(Ome)-fmk (#FK009), z-VAD-fmk (#FK109), Q-VD-OPh (#03OPH109-CF), were purchased from MP Biomedical (Santa Ana, USA). Q-VE-OPh (#A0007) was purchased from Apoptrol LLC (Beavercreek, Ohio, USA) and Emricasan (#510230), also known as IDN 6556 and PF 03491390, was obtained from Medkoo Biosciences (Morrisville, NC, USA). TRP601 (C_40_H_48_F_2_N_6_O_11_; MW: 826.84) and Δ2Me-TRP601 (C_38_H_44_F_2_N_6_O_11_; MW: 798.79) were provided by Chiesi Parmaceuticals [31]. TRP801 (Q-LETD(OMe)-OPh) (MW: 771.76) was custom-synthetized by Polypeptides Laboratories (Strasbourg, France) [31]. LJ2 was purchased (custom-synthesis) from by Polypeptides Laboratories (Strasbourg, France). LJ2a, LJ2b, LJ3a, and LJ3b were purchased (custom synthesis) from Bachem Americas, Inc. (Vista, CA, USA). Caspase inhibitors were solubilized in DMSO at 10 mM and stored at −80°C.

### In vitro Caspases activity and Inhibition assays

Caspase activities were determined by monitoring the hydrolysis of appropriate fluorogenic substrates, in 96 wells plates using a BMG Fluostar microplate reader, as a function of time at 37 °C in the presence of untreated caspase (control) or enzyme that had been incubated with a test compound. Substrates and compounds were previously dissolved in DMSO at 10 mM, with final solvent concentration kept constant at lower 4% (v/v). Initial velocity (V0) was determined from the linear portion of the progress curve. The composition of the activity buffer was: 20 mM HEPES, pH 7.4; 0,1 %CHAPS, 5 mM DTT, 2 mM EDTA; 800 mM sodium succinate for Caspase-2 (0.2 nM) with 25 μM of Ac-VDVAD-Amino-methylcoumarin (AMC) and 20 mM HEPES, pH 7.4; 0,1 %CHAPS, 5 mM DTT, 2 mM EDTA; supplemented with 1 mg/mL of stabilizing agent BSA for Caspase-3 (0.1 nM) with 10 μM of Ac-DEVD-AMC. Compounds (0.001 – 100 μM) were tested in duplicate for each inhibitor concentration to detect their inhibitory potential. Enzyme and compounds were incubated for 30 min before to initiate enzymatic reaction by adding substrate. Initial rates determined in control experiments (V0) were considered as 100 % of the caspase activity; initial rates that were below 100 % in the presence of tested compound (Vi) were considered as inhibition. The inhibitory activity of compounds was expressed as IC_50_ (Inhibitor concentration giving 50 % inhibition). The values of IC_50_ were calculated by fitting the experimental data using Mars data Analysis 2.0 and KaleidaGraph softwares.

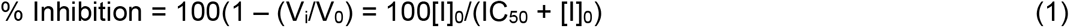

### Enzyme Mechanistic Studies

For noncovalent (reversible) inhibitors, the mechanisms of inhibition were determined by varying substrate and inhibitor concentrations and using Lineweaver-Burk representations.

For suicide inhibitors, inactivation can be represented by the minimum kinetic scheme (eq 2), where E and I are the free forms of enzyme and inhibitor, E*I a kinetic chimera of the Michaelis complex and E-I the covalent complex or inactivated enzyme.

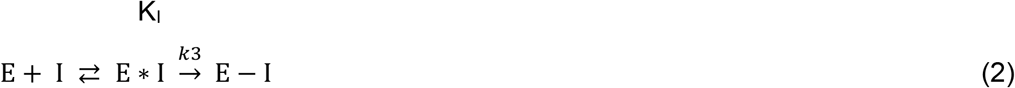

K_1_ and k_3_ are the kinetic constants characterizing the inactivation process and the k_3_/K_I_ ratio is an index of the inhibitory potency. First-order rate constant, k_3_, and dissociation constant, K_I_, were determined for Caspase-2 et −3 using the progress curve method [35]. Briefly, the enzyme activities were measured continuously for 120 min. To determine the kinetic parameter, progress curves were obtained at several inhibition concentrations using fixed substrate concentration. Product released for each inhibitor concentrations was plotted versus time following eq 3 where the constant *π* depended on [I]’, a modified inhibitor concentration due to substrate competition, according to eq 4. [I]’ is defined by eq 5, where K_M_ was Michaelis constant for the enzymatic hydrolysis of the appropriate fluorogenic substrate. The ratio k_i_/K_I_ was obtained by fitting the experimental data to the equations (F.U., fluorescence unit):

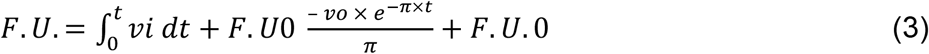

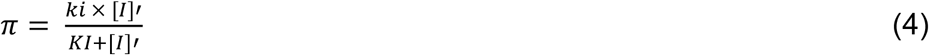

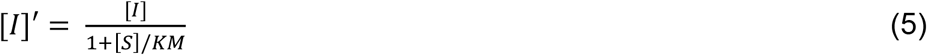

Linear and nonlinear regression fits of the experimental data to the equations were performed with KaleidaGraph software. The experimental conditions were [Caspase-2]_0_ = 0.2 nM, [Ac-VDVAD-AMC]_0_ = 25 μM and [Caspase-3]_0_ = 0.1 nM, [Ac-DEVD-AMC]_0_ = 10 μM; [I]_0_ = 0 – 100 μM.

### Cell lines and cell death modulation

HeLa cells (obtained from ATCC) were cultured in Dulbecco’s Modified Eagle Medium (DMEM, High Glucose, GlutaMAX™, Pyruvate) supplemented with antibiotics and 10% FCS (Gibco, Life technologies) and maintained in T75 Flasks at 1×10^6^ cells per flask. 24h before treatments HeLa cells were transferred to 6-well plates (8×10^4^ cells/well). The lymphoblastoid cell line Molt4cl8 was a gift from Dr Bernard Krust (Inserm, University Paris 5) and was cultured in RPMI 1640 Glutamax medium supplemented with Hepes, antibiotics, and 10% FCS. HEK293 cells were obtained from ATCC (CRL-1573) and cultured in DMEM with Glutamax and 25 mM glucose (Life Technologies) supplemented with 10% fetal calf serum (FCS) and antibiotics. For cell death induction, the following agents were added to cell cultures: Vincristine (Sigma) and Panobistat (LBH589; Selleck). Cell death was analyzed using fluorescence microscopy (Axio-observer Z1, Zeiss; equipped with CCD camera CoolsnapHQ2, Ropert Scientific) and cytofluorometry (MACSQuant® VYB; Miltenyi Biotec, Bergisch Gladbach, Germany) after labelling with propidium iodide (1μg/mL; 10 min; 37°C; Life Technologies) and Hoechst 33342 (1μg/mL; 5 min; RT; Sigma).

### β-Amyloid preparation

Lyophilized and HPLC-purified Human β-Amyloid_1-42_ (Aβ_1-42_) peptide was purchased from Tocris (#1428). 1,1,1,3,3,3-Hexafluoro-2-propanol (HFIP) was purchased from Sigma Aldrich (Germany). Oligomeric and non-oligomeric form of Aβ_1-42_ peptide were produced according to ref [36]. Briefly, lyophilized peptides were solubilized at 1 mM in 1,1,1,3,3,3-Hexafluoro-2-propanol (HFIP). After 30 min of incubation at room temperature, HFIP was evaporated overnight and peptides were dried (Speed Vac, 1h 4°C). Then, Aβ peptide stock solution was obtained by solubilization at 5 mM in DMSO followed by bath sonication for 10 min. To obtain oligomers, Aβ_1-42_ stock solution was diluted to 100 μM in phenol free DMEM-F12 medium and then incubated at 4°C, 24h. Non-oligomeric form was obtained by diluting Aβ_1-42_ stock solution in fresh milliQ water.

### Fluidic microsystems

Microfluidic chips were produced by standard molding methods [37] using epoxy-based negative photoresists (SU-8) and MicroBrain Biotech proprietary microdesigns (Brainies™, Cat#: MBBT4 and MBBT5; Marly le Roi, France). Briefly, Polydimethylsiloxane (Sylgard 184, PDMS; Dow Corning, MI, USA) was mixed with curing agent (9:1 ratio) and degassed under vacuum. The resulting preparation was poured onto a chosen SU8 mold and reticulated at 70°C for 2 at least hours. The elastomeric polymer print was detached, and 2 reservoirs were punched for each chamber. The polymer print and a glass cover slip were cleaned with isopropanol, dried, and treated for 3 minutes in an air plasma generator (98% power, 0.6 mBar, Diener Electronic, Ebhausen, Germany) and bonded together. The day before neuronal seeding, chips were UV-sterilized for 20 min, then coated with a solution of poly-D-lysine (10 μg.ml^−1^ Sigma #P7280, St. Louis, MO, USA), incubated overnight (37°C, 5% CO_2_), and rinsed 3 times with Dulbecco’s phosphate buffer saline (D-PBS) (Thermo Fisher Scientific, Invitrogen #14190169, Waltham, MA, USA). Then, 4 hours before cell seeding, chips were treated with a solution of Laminine (5 μg/mL: Sigma) in D-PBS. Brainies™MBBT5 is a chip with a design containing 4 neuronal diodes. One neuronal diode includes 2 rectangular culture chambers (volume ~1 μL) each connected to 2 reservoirs and separated by a series of 500 μm-long asymmetrical micro-channels (3 μm high, tapering from 15 μm to 3 μm). Brainies™MBBT4 is a chip containing 8 rectangular culture chambers (volume ~1 μL) each connected to two reservoirs but devoid of microchannels to connect chambers.

### Primary neuron microcultures

All animals were ethically maintained and used compliance with the European Policy and Ethics. E16 embryos (Swiss mice, Janvier, Le Genest Saint Isle, France) were micro-dissected in Gey’s Balanced Salt Solution (GBSS, Sigma (without CaCl_2_ and MgCl_2_) supplemented with 0,1% (w/v) glucose (Invitrogen). Structures were digested with papain (20U/mL, Sigma #76220) in DMEM Glutamax (31966; Invitrogen) for 15 min at 37°C. After papain inactivation with 10 % (v/v) of fetal bovine serum (GE Healthcare, U. K.) structures were mechanically dissociated in DMEM Glutamax containing DNAse-I (100 U/mL, D5025, Sigma). After 10 min centrifugation at 700 *g*, cortical and hippocampi cells were resuspended in DMEM Glutamax supplemented with 10 % FBS, 1% streptomycin/penicillin (Life Technologies), N2 supplement (17502048; Thermo Fisher Scientific), and B-27 supplement (17504-044, Thermofisher Scientific). For the monoculture model (MBBT4 devices), 20.10^3^ hippocampal neurons were seeded in each chamber. For the compartmentalized coculture model (neuronal diode in MBBT5 devices), 40.10^3^ cortical cell and 15.10^3^ hippocampal cells were seeded in each input chambers and output chamber, respectively. The culture medium was renewed every 5 days. Microfluidic chips were placed in Petri dishes containing 0,1 % EDTA (Sigma) and incubated at 37°C in 5% CO_2_ atmosphere. In the neuronal diodes, cortical axons entered the microchannels and reached the hippocampi chamber in around 4-5 days. In both the monoculture and coculture systems, cells were cultured for 3 weeks to allow high dendrites and synapses density and intense electrical firing. GFAP staining, consistently showed that astrocytes represented less than 5% of the cultures.

### β-Amyloid and pharmacological treatments in microculture models

Cells were treated with caspase inhibitors and/or Aβ_1-42_ at 20 days *in vitro* (DIV). For the hippocampal monocultures, the culture medium was removed and replaced by fresh medium optionally containing 10 nM Aβ_1-42_ oligomers or monomers, with or without caspase inhibitors. After 6h, cells were fixed, permeabilized, and subjected to immunolabelling. For cortico-hippocampal compartmentalized cocultures, the medium of the input chamber (cortical) was replaced by fresh medium optionally containing 100 nM Aβ_1-42_ oligomers or monomers, whereas the medium of the output (hippocampal) chamber was replaced by fresh medium optionally containing, or not, caspases inhibitors. To ensure fluidic isolation between input and output chamber of each neuronal diodes, a differential hydrostatic pressure between chambers was maintained. After 6h, cells were fixed, permeabilized, and hippocampal neurons (output compartment) were subjected to immunolabelling.

### Immunofluorescence, detection, and image acquisition

Immunostaining was performed directly in microfluidic chambers as described [38]. Briefly, neurons were fixed for 20 min at RT in D-PBS containing 4% (w/v) paraformaldehyde (PFA) (Euromedex #15714-S, Souffelweyersheim, France) and 4% (w/v) sucrose (Sigma #S0389). Cells were then washed once with D-PBS for 10 min and permeabilized for 30 minutes with D-PBS containing 0,2% (v/v) Triton X-100 (Sigma) and 1 % (w/v) BSA (Sigma). Solutions of primary antibodies, diluted in PBS, were incubated for 2h at RT. Cultures were rinsed 2 times for 5 min with PBS and further incubated with the corresponding secondary antibodies for 2 h at room temperature. For characterization of neuronal culture quality, the following conjugated antibodies were used: anti-βIII tubulin-Alexa Fluor 488 (1:500, AB15708A4; Millipore), anti-microtubule-associated protein 2 (MAP2) Alexa 555 (1:500, MAB3418A5; Millipore), antimicrotubule-associated protein 2 (MAP2) Alexa Fluor 647 (1:500, NB120-11267AF647; Millipore). Cell nuclei were stained by using Hoescht 33342 (2 μg/mL, Sigma). For synapse loss studies, dendritic spine actin-F was labelled with Phalloidin Alexa Fluor 555 (1:100, A34055; Thermo Fisher Scientific) and the following antibodies were used anti-Bassoon (1:400, SAP7F407; mouse monoclonal; Enzo Life) and anti α-synuclein (1:500, D37A6; rabbit polyclonal; Cell Signaling). Species-specific secondary antibodies coupled to Alexa 488 and 350 (1:500; Invitrogen) were used. Images were acquired using an Axio-observer Z1 microscope (Zeiss, Wetzlar, Germany) fitted with a cooled CCD camera (CoolsnapHQ2, Ropert Scientific, Trenton, NJ, USA). During acquisition, the microscope was controlled with MetaMorph® Microscopy Automation & Image Analysis Software. Images were analyzed using ImageJ software (NIH, Bethesda, MD, USA).

### Quantification of Synaptic Disconnection

Synaptic disconnection was assessed as described [39] with slight modifications. Briefly, in hippocampal monocultures, synaptic disconnection was assessed through fluorescence microscopy by counting phalloidin clusters affixed to MAP2 and Bassoon. In cortico-hippocampal compartmentalized cocultures, synaptic disconnection was assessed by counting α-Synuclein presynaptic clusters affixed to MAP2 positive hippocampal dendrites. All images were obtained using the same acquisition parameters. The images were similarly processed with ImageJ software before being used for quantification: the brightness/contrast of all control images was optimized manually to eliminate the background and to maximize the signal. The means of the minimum and maximum intensities were then calculated in the control condition and these settings were applied to all images. The brightness / contrast of all images was optimized manually to eliminate the background and to maximize the intensity of the signal. α-synuclein/ MAP2 and Phalloidin/MAP2/Bassoon merges were then used for quantification. The number of spines was determined by counting individual clusters along 10 dendritic region (100 um length each) in three independent experiments for every epitope. Only synapses detected on or near the neurites were included for analysis. The resulting synapse counts were then exported to Excel for further analysis. Reported values are means for at least three independent experiments, each performed in triplicate.

### Transfections and Immunoblot analysis

To test SREBP processing, HEK 293 cells were plated at a density of 3 × 10^6^ per well of 10 cm plates. The next day, 3 μg of indicated cDNAs were transfected using lipofectamine 3000 (Thermo Fisher Scientific, MA) according to manufacturer’s instruction. Whole cell lysates (WCL) were prepared in lysate buffer (150 mM Tris-HCl, pH 7.4, 10% sodium-deoxycholate, 100 mM NaCl, 100 mM EDTA, 100 mM PMSF, 200 mM NaF, 100 mM Na_3_VO_4_, and a mixture of protease inhibitors. Equal amounts of WCL were subjected to immunoblot (WB) analysis with anti-V5 monoclonal antibody (Cell Signaling Technology; Cat#13202) [12].

### NMR experiments

Nuclear Magnetic Resonance (NMR) experiments were carried out on a 600 MHz spectrometer equipped with a cryoprobe. LJ3a and LJ3b samples were solubilized in 100% deuterated DMSO and ^1^H NMR experiments (1D ^1^H, 2D TOCSY [Suppl Ref 1], 2D COSY) [Suppl Ref 2], [Suppl Ref 3], 2D NOESY [Suppl Ref 4], [Suppl Ref 5]) (Supplementary material, Figure S1A and Figure S2A), ^13^C experiments in natural abundance (1D ^13^C, 2D ^1^H-^13^C HSQC [Suppl Ref 6], 2D ^1^H-^13^C HMBC) [Suppl Ref 7]) (data not shown) and ^15^N experiments in natural abundance ^1^H-^15^N SOFAST-HMQC [Suppl Ref 8] (Fig. 6a,b) were recorded. The ^1^H-^15^N SOFAST-HMQC [Suppl Ref 8] experiments in natural abundance allow to register a twodimensional heteronuclear correlation spectrum of the pseudo-peptide and make it possible to correlate the chemical shift of one amide proton on the abscissa axis with the chemical shift of the nitrogen which carries it on the ordinate axis for each of the amino acids (Fig. 6a, b). The ^1^H TOCSY [Suppl Ref 1] experiment makes it possible to identify all the protons belonging to the same spin system thanks to the magnetization transfer through the bonds. Then, the succession and the order of the different residues in the pseudo-peptide is identified thanks to the NOESY [Suppl Ref 4], [Suppl Ref 5] experiment which will make it possible to connect the different residues thanks to a transfer of magnetization through space [Suppl. Fig. S1A, S2A].

**Figure 1:**
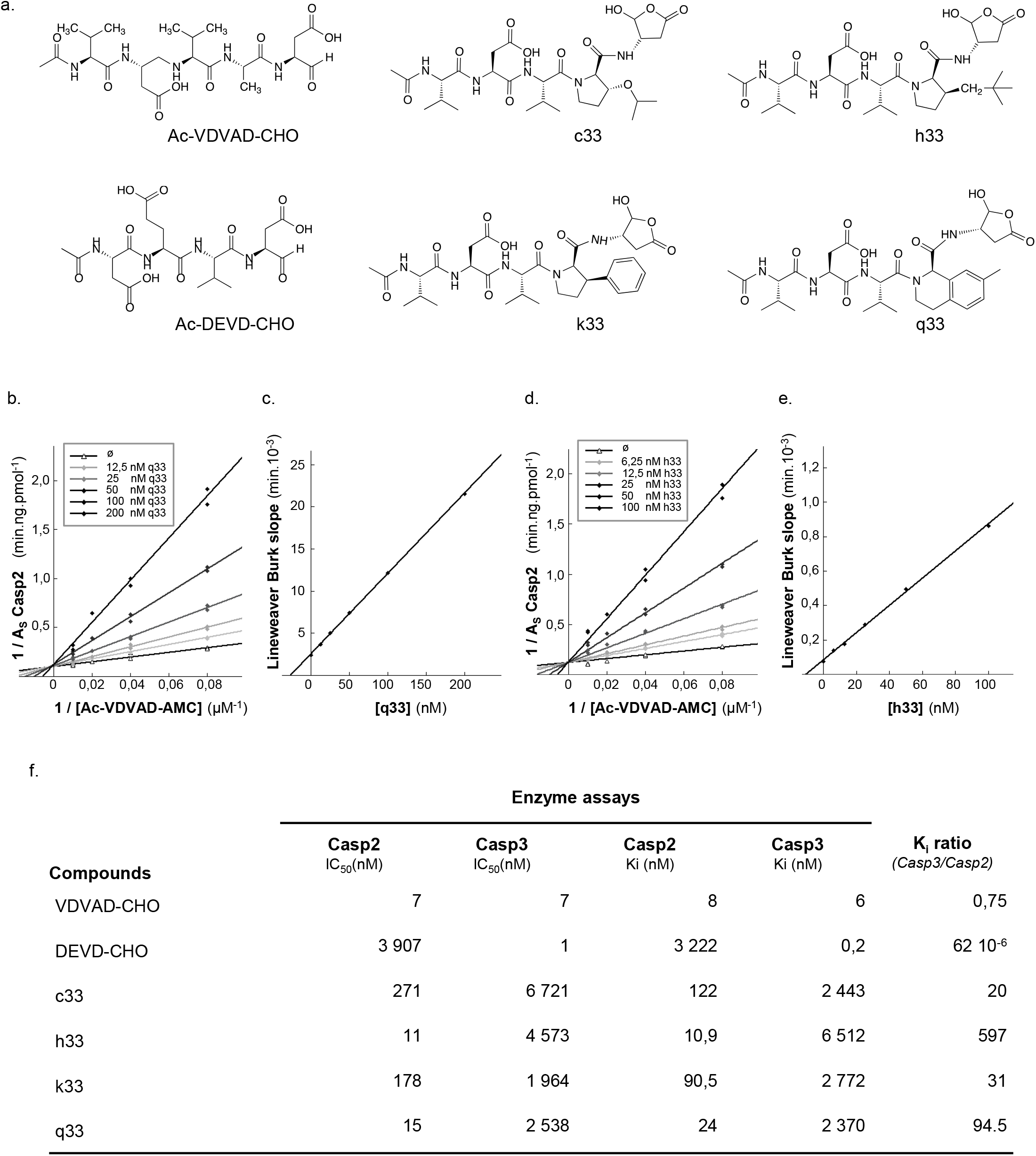
Kinetic parameters on Caspase-2 versus Caspase-3 of P2-modified VDVAD pentapeptides derivatives. **a**) Structures of reversible Caspase-2/3 inhibitors. Ac-VDVAD-CHO is a non-selective canonical pentapeptide Caspase-2/3 inhibitor, Ac-DEVD-CHO is a canonical tetrapeptide Caspase-3 inhibitor. c33, k33, q33, h33 are VDVAD derivatives where the P2 Alanine residue was replaced by bulky substituent to decrease activity against Caspase-3. **b-e**) Potent competitive inhibition of Caspase-2 by q33 (**b, c**) and h33 (**d, e**). Lineweaver-Burk plots of Casp2 inhibition with q33 (**b**) and h33 (**d**) are shown together with the relationship between the slopes of lines in Lineweaver-Burk plot analyses and the concentration of q33 (**c**) and h33 (**e**) inhibitors. **f**). IC50, Ki values and Casp3/Casp2 selectivity ratio of reversible P2-modified VDVAD derivatives. Data points on the graphs represent the mean ± SD, and calculated *K*_i_ values represent mean ± standard error (SE) from 3 independent experiments.

**Figure 2:**
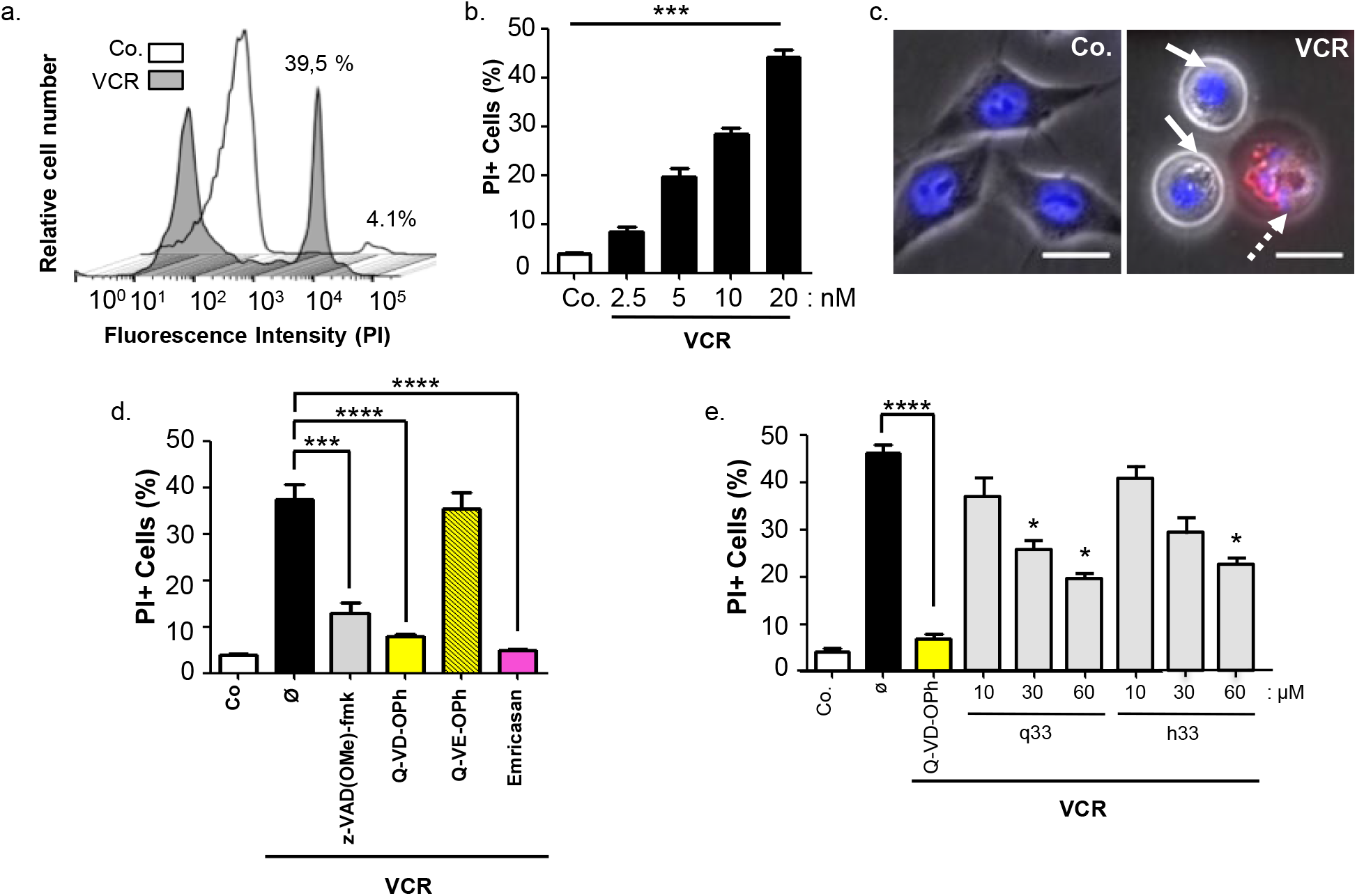
q33 and h33 are protective against the cytocidal effect of Vincristine. **a)** Vincristine induces HeLa cells death. Cells were incubated for 44 h at 37°C with 20 nM Vincristine (VCR), then incubated with propidium iodide at 2 μg/mL for 10 min and analyzed by flow cytometry to quantify plasma membrane permeabilization. Histograms show fluorescence intensity detected by flow cytometry (one representative experiment) in the presence (VCR) or Absence (Co.) of Vincristine. % of propidium iodide positive cells are indicated for both conditions. **b)** Dose-response of Vincristine-induced cell death. HeLa cells were incubated and treated as in (a) with the indicated doses of Vincristine (VCR) and analyzed by flow cytometry. Histogram show the mean ±SD of 5 independent experiments (***p value < 0.001). **c**) Vincristine induces apoptotic nuclear morphology in HeLa cells. Representative fluorescence microscopy micrographs of cells treated for 44 h with 20 nM vincristine (VCR) or not (Co.) and stained with Hoechst 33342. Vincristine induces progressive nuclear changes with first condensed nuclei (arrows) and then apoptotic bodies (dashed arrow). **d**) Broad-spectrum irreversible caspase inhibitors prevent vincristine-induced cell death. HeLa cells were pretreated for 1hr with the indicated broad-spectrum irreversible caspases inhibitors (z-VAD(OMe)-fmk at 30 μM, Q-VD-OPh 30 μM, Emricasan 10 μM, or the negative control Q-VE-OPh 30 μM), then treated with vincristine as in panel a, and analyzed by flow cytometry. Histograms show the mean +/-SD of 3 independent experiments (***p value < 0.001). **e**) Effect of the reversible Caspase-2 inhibitors q33 and h33 against vincristine-induced cell death. HeLa cells were pretreated for 1hr with the indicated doses of q33 or h33, then treated with vincristine as in panel a, and analyzed by flow cytometry. Histograms show the mean +/-SD of 3 independent experiments (*p value < 0.1).

**Figure 3:**
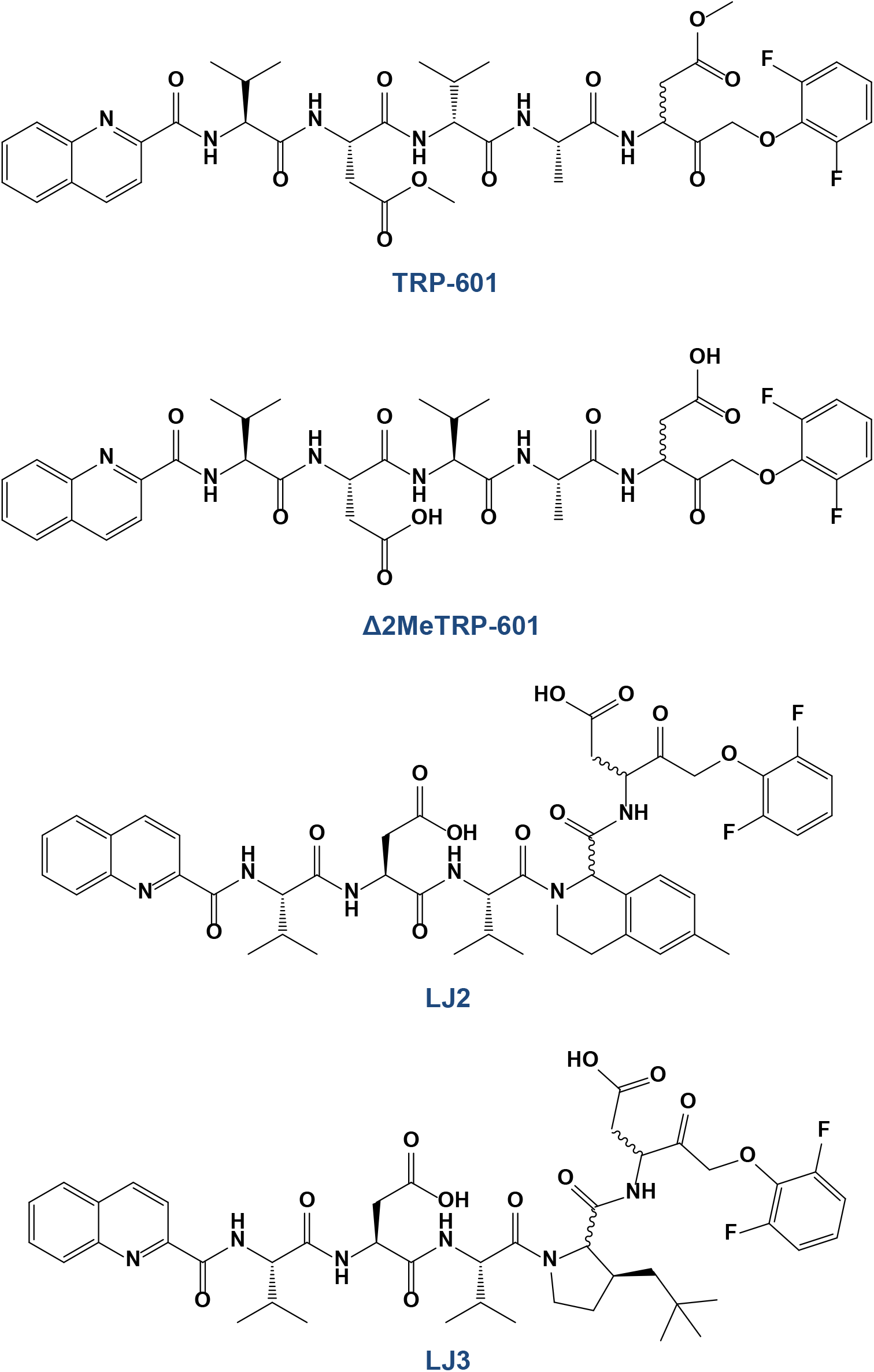
structure of Caspase-2 irreversible inhibitors. Quinolin-2-carbonyl-VD(OMe)VAD(OMe)-CH_2_-O(2,6F_2_)Ph (referred as TRP601 [31]) and Quinolin-2-carbonyl-VDVAD-CH_2_-O(2,6F_2_)Ph (referred as Δ2Me-TRP601) are pentapeptide derivatives. Quinolin-2-carbonyl-VDV-(methyl-isoquinolyl)-D-CH_2_-O(2,6F_2_)Ph (referred as LJ2) and Quinolin-2-carbonyl-VDV(3-neopentyl)D-CH_2_-O(2,6F_2_)Ph (referred as LJ3), are pentapeptide-derived peptidomimetics.

**Figure 4:**
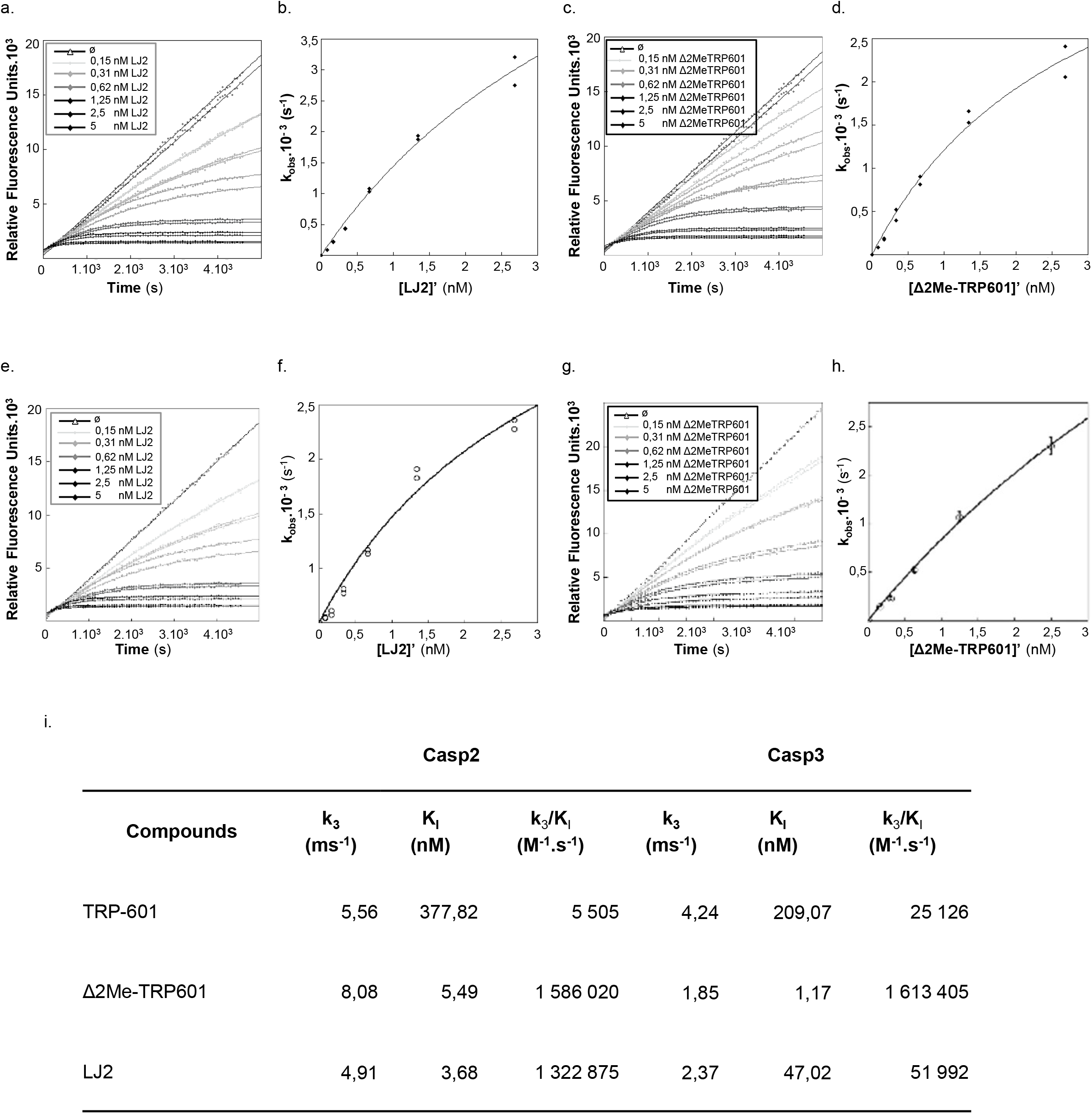
kinetic of Caspase-2 vs Capsase-3 inhibitions by TRP601, Δ2Me-TRP601, and LJ2. **a, c**. Progress curves of Ac-VDVAD-AMC hydrolysis catalyzed by caspase-2 in the presence of various concentrations of LJ2 (a) and Δ2Me-TRP601 (c), as indicated. The data were obtained under the conditions described in the Materials and methods section (25 μM substrate and 0.2 nM Caspase-2) and analyzed according to Eq. (2–5). **b**. Values of *k*_obs_ obtained from panel “**a**” plotted versus LJ2 concentrations. **d**. Values of *k*_obs_ obtained from panel “**c**” plotted versus Δ2Me-TRP601 concentrations. **e, g**. Progress curves of Ac-DEVD-AMC hydrolysis catalyzed by caspase-3 in the presence of various concentrations of LJ2 (e) and Δ2Me-TRP601 (**g**), as indicated. The data were obtained under the conditions described in the Materials and methods section (10 μM substrate and 0.1 nM Caspase-3) and analyzed according to Eq. (2). **f**. Values of *k*_obs_ obtained from panel “**e**” plotted versus LJ2 concentrations. **h**. Values of *k*_obs_ obtained from panel “**g**” plotted versus Δ2Me-TRP601 concentrations. **i.** The plots in **b**, **d**, **f**, and **h** were fit according to Eq. (3) to generate K_I_ and *k*_inact_ (k_3_) respective values. Indicated K_i_ and k_3_ value of each inhibitor on Caspase-2 and Caspase-3 are mean values from five independent experiments (SE < 0.07). Koff rates (k_3_/K_i_) and Casp3/Casp2 selectivity ratio (K_off_ rates Casp3 / K_off_ rates Casp2) are also indicated.

**Figure 5:**
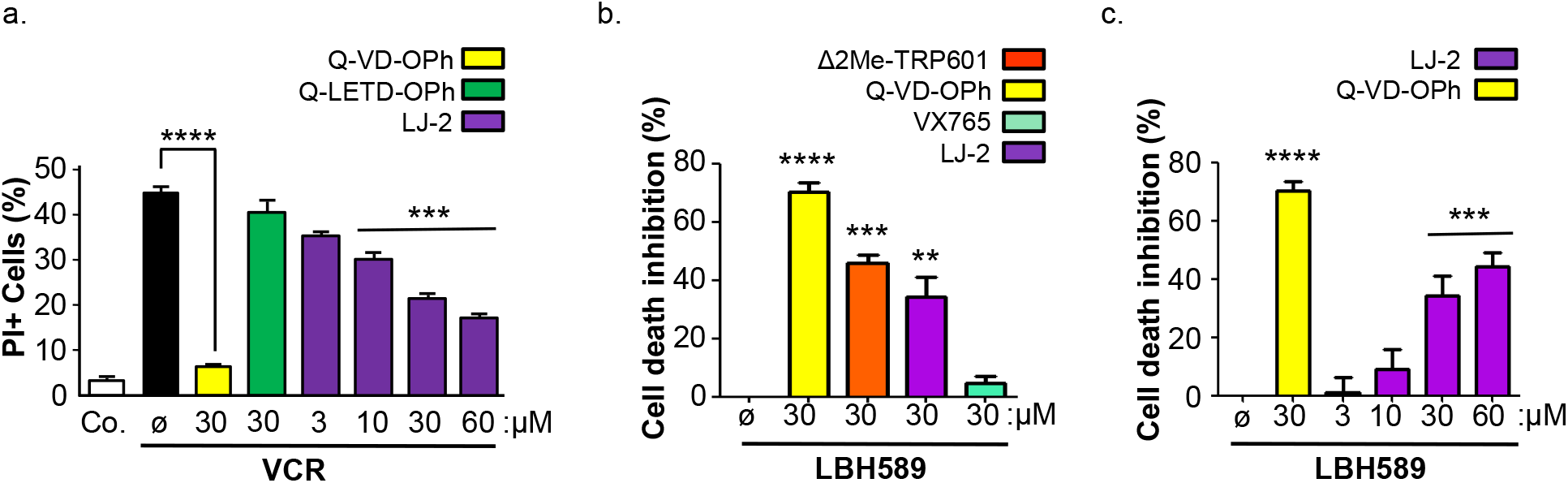
LJ2 inhibits cell death induced by microtubule destabilization and hydroxamic acid-based deacetylase (HDAC) inhibition. **a)** Effect of the irreversible Caspase-2 inhibitors LJ2 against vincristine-induced cell death. HeLa cells were pretreated for 1hr with the indicated doses of LJ2, Q-VD-OPh, or Q-LETD-OPh, then treated with vincristine (VCR) as in Figure 2, then incubated with propidium iodide at 2 μg/mL for 10 min and analyzed by flow cytometry. Histogram data are mean +/-SD (n=3, *p value < 0.01). **b)** Effect of Caspases-inhibitors against Panobistat (LBH589)-induced cell death. Molt4 lymphoid human cells where were pretreated for 1hr with 30 μM of LJ2, Q-VD-OPh, or Belnacasan (VX-765), then treated for 24h with 25 nM of LBH589, then incubated with propidium iodide as in (**a**) and analyzed by flow cytometry. Histograms data are mean +/-SD (n=4, ***p value < 0.001). **c)** Dose-response of LJ2-mediated cell death inhibition. Molt4 cells were treated as in (**b**) with the indicated doses of LJ2 or Q-VD-OPh and analyzed by flow cytometry. Histogram data are mean +/-SD (n=3, **p value < 0.01).

**Figure 6:**
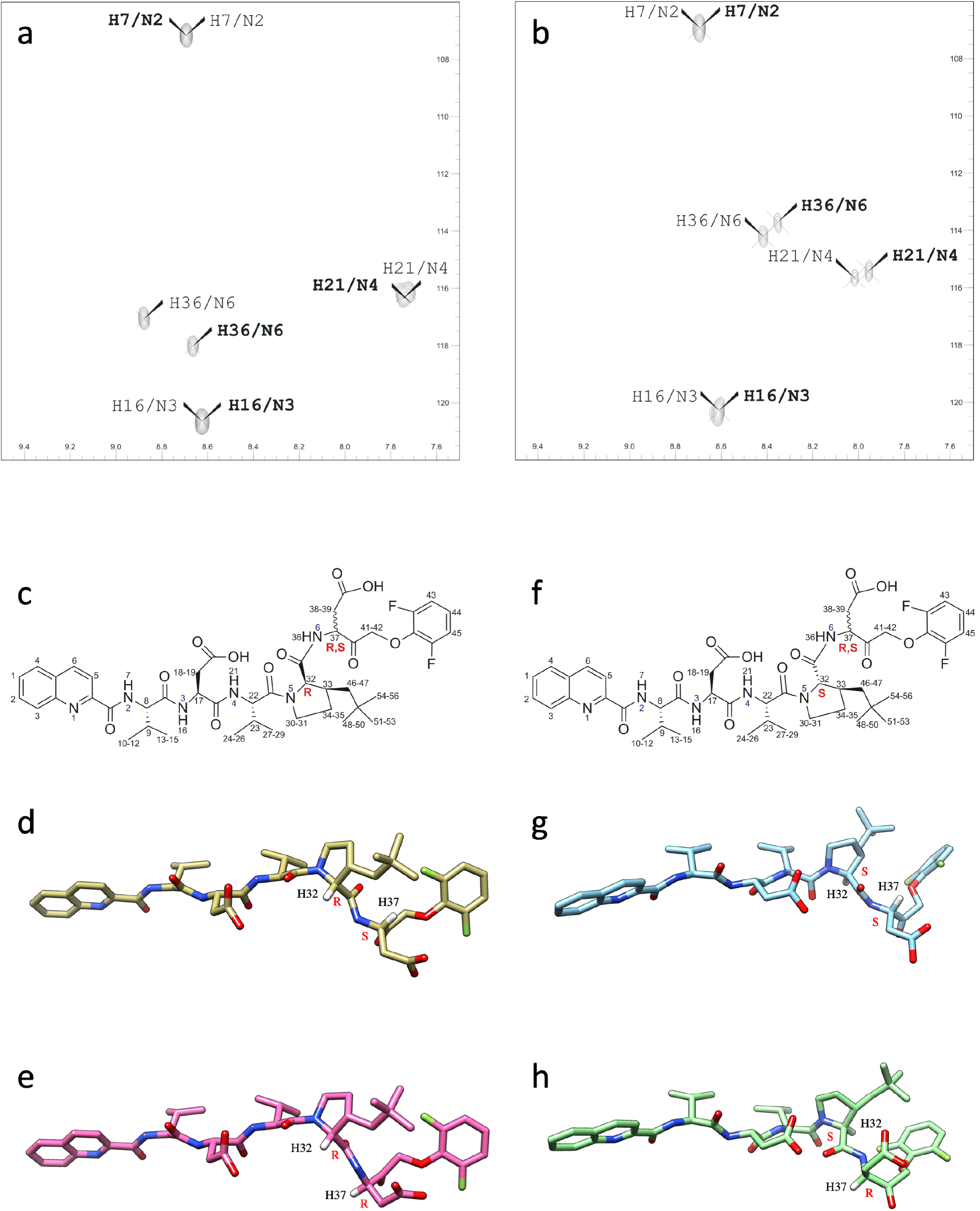
determination of the absolute configuration of LJ3a and LJ3b by NMR spectroscopy. **a.** Heteronuclear Multiple Quantum Coherence (HMQC) for LJ3a. Two-dimensional (2D) ^1^H-^15^N 2D heteronuclear HMQC spectrum of the LJ3a sample showing the correlation between the nitrogen and the proton it carries. The spectrum shows additional peaks due to the racemization of P1 and corresponding to two different isomers. The assignment has been made and the correlations for one isomer are represented in thin letters while the other is in bold. **b.** HMQC for LJ3b. 2D ^1^H-^15^N 2D heteronuclear HMQC spectrum of the LJ3b sample showing the correlation between the nitrogen and the proton it carries. The spectrum shows additional peaks due to the racemization of P1 and corresponding to two different isomers. The assignment has been made and the correlations for one isomer are represented in thin letters while the other is in bold. **c.** LJ3a molecules of general formula Qco-P5-P4-P3-P2-P1-OPh with racemization in P1 (Asp-OPh) and absolute configuration R in position P2 (3-neopentyl proline). The positions P5 (Val 5), P4 (Asp 4) and P3 (Val 3) have an absolute configuration S. **d.** Structural 3D model of the 1S2R isomer. **e.** Structural 3D model of the 1R2R isomer. **f.** LJ3b molecules of general formula Qco-P5-P4-P3-P2-P1-Oph with racemization in P1 (Asp-OPh) and absolute configuration S in position P2 (3-neopentyl proline). The positions P5 (Val 5), P4 (Asp 4) and P3 (Val 3) have an absolute configuration S. **g.** Structural 3D model of the 1S2S isomer. **h.** Structural 3D model of the 1R2S isomer.

### Theoretical model of the 3D structures

The theoretical 3D structure of each LJ3 isomers (1R, 2R), (1S, 2R), (1R, 2S) and (1S, 2S) was built and a combination of steepest descent and conjugate gradient methods was used to minimize these structures using UCSF Chimera software 1.12 [Suppl Ref 9]. The distances between protons specific to one or the other isomer were measured on the resulting structures and were compared to the volumes of the NOEs measured on the NOESY spectra of each isomer and to the corresponding NOEs derived distances [Suppl. Methods] [Suppl. Fig. S1, S2].

### Statistical analysis

Data are represented as mean ± SEM or ± SD as indicated. Differences in mean values were analyzed by Student t-test or one-way ANOVA (for more than 2 groups) and post-hoc Tukey test. For all analysis **** p < 0.0001; *** p < 0.001; ** p < 0.01; * p < 0.1. Statistical analyses were performed using GraphPad Prism 9 software.

## Results

### S_2_ pocket of the Caspase-2 active site as a determinant for inhibitor selectivity

The S2 subsite of Casp2 forms a long groove that runs perpendicular to the active site cleft, whereas the Casp3 S2 subsite is lipophilic and forms a round, bowl-like shape that can preferentially bind shorter hydrophobic resides.^32^ Targeting the Casp2 S2 subsite with either longer R-groups or bulkier amino acids (that would likely sterically clash with Tyr204 in Casp3) could be an effective strategy in selectively targeting Casp-2. As a starting point, we have evaluated, on both recombinant Casp2 and Casp3, the IC_50_ and K_i_ values of a series of pentapeptide-based reversible Casp-2 inhibitors including the canonic Ac-VDVAD-CHO, and derivatives modified at the P2 position. The four P2-modified pentapeptides aldehydes (Fig.1a), namely c33, k33, q33, and h33, where chosen from a medicinal chemistry series reported by Maillard et al, because these authors showed IC_50_ values indicating improved selectivity toward Casp2 (reduced Casp3 inhibition) as compared to Ac-VDVAD-CHO [32]. We extended the characterization of these inhibitors and found that q33 and h33 present K_i_ ratio (Casp3/Casp2) of 94,5 and 597, respectively (Fig.1b-f). When compared to the Ac-VDVAD-CHO K_i_ ratio (Casp3/Casp2), this represents 126 times and 796 times improvement of selectivity for q33 and h33, respectively.

As caspase-2 is reportedly required for cell death induced by cytoskeletal disruption [40], we used the depolymerizing agent vincristine as a cell killing model to evaluate the cytoprotective effect of q33 and h33. Vincristine killed HeLa cells at nanomolar concentrations (Fig. 2a, b), and dying cells showed typical nuclear apoptotic morphology (Fig.2c; Hoechst 33342, blue). In this model, cell death is strictly Caspase-dependent, as 30 μM of the standard broad-spectrum caspase inhibitors z-VAD(Ome)-fmk and Q-VD-OPh, but not its control Q-VE-OPh, inhibited more than 80% and 90% of cell death, respectively (Fig.2d). This was further confirmed with 10μM of Emricasan, a highly potent pan-caspase inhibitor, which abolished 95% of Vincristin-mediated cell death (Fig.2d). Furthermore, the reversible Casp2 inhibitors q33 and h33 showed cytoprotective effect against vincristine-induced killing (Fig.2e). However, the relation dose-effect was not significant, possibly due to the kinetics characteristics of the Vincristin killing and to the reversible properties of h33 and q33.

### Design and enzyme kinetics of Casp2-selective irreversible peptidomimetics

Consequently, we reasoned that a potent and selective Caspase-2 inhibitor should combine the Caspase-2 selectivity of q33 or h33 with the bioavailability, irreversible inhibition, and efficiency of the previously developed group-II caspase inhibitor Q-V(Ome)DVA(Ome)D-OPh (also known as TRP-601), or its highly active metabolite Q-VDVAD-OPh (also known as Δ2Me-TRP601) [31]. We consequently designed LJ2 and LJ3, two peptidomimetics of general structure Q-Val-Asp-Val-X-Asp-OPh (Q-VDVXD-OPh) where Q is a quinaldoyl, OPh is a 2,6-difluorophenyloxymethylketone, and X is either a 6-methyl-1,2,3,4-tetrahydroisoquinoline-1-carbonyl (within LJ2) or a 3-neopentyl-Proline (within LJ3) (Fig. 3).

The inhibitory potency of the newly synthetized LJ2 compound toward recombinant human Casp2 was evaluated. LJ2, as Δ2Me-TRP601, behaved as a dose- and time-dependent inhibitor of Casp2 (Fig4a-d; LJ2/Casp2 IC_50_ = 0.37 ±0,01 nM; Δ2Me-TRP601/Casp2 IC_50_ = 0.53±0,02 nM; n=5). Δ2Me-TRP601 confirmed to be a powerful inhibitor of Casp3 (Δ2Me-TRP601/Casp3 IC_50_ = 0.31±0,01 nM; n=5) [31], whereas LJ2 had a 350 times lower effect on Casp3 as compared to Casp2 (LJ2/Casp3 IC_50_ of 140.3 ±5 nM; n=5) (Fig.4e-h). Then, the kinetic parameters k_3_/K_i_, k_3_, and K_i_, were determined (Fig.4i). LJ2 and Δ2Me-TRP601 are both excellent inactivator of Casp2 showing k_3_/K_i_, ratio of ~1 322 875 M^−1^s^−1^ and ~1 586 020 M^−1^s^−1^, respectively. LJ2 was still able to inactivate Casp3 but 25 times less than Casp2 (LJ2/Casp3 k_3_/K_i_, ratio = 51 992 M^−1^s^−1^; n=3).

### LJ2 inhibits cell death in human cell lines subjected to microtubule destabilization or histone deacetylase inhibition

We then evaluated the cytoprotective potential of LJ2 in cell death assays. When added to cell lines or primary neurons in culture, LJ2 was devoid of any toxicity at least up to 100 μM (not shown). LJ2 significantly prevented Vincristin-induced cell death in a dose-dependent manner (Fig 5a). A similar cytoprotective pattern was found with the group-II caspase inhibitor Δ2Me-TRP601, whereas the Caspase-8 inhibitor Q-LETD-OPh (also known as TRP801) had no effect on Vincristin-induced cell killing. We also evaluated the effect of LJ2 against cell death induced by the deacetylase inhibitor Panobistat (LBH589) in lymphoid human T cells lines. Reportedly LBH589 induces caspase-2 dependent apoptotic cell death in lymphoid cell lines [41]. Accordingly, LJ2 showed a dose-dependent inhibition of cell death (Fig. 5b). In this model the pan Caspase inhibitor Q-VD-OPh, strongly inhibited cell death, whereas inhibition of the inflammatory caspases (Casp1, −4, −5) with Belnacasan (also known as VX-765) did not affect cell death (Fig. 5b,c).

### Optimization of highly effective and potent Casp2-selective inhibitors

Synthesis of LJ2 leads to racemization in P2 and P1. Consequently, the compound LJ2 is a mixture of 4 diastereoisomers. Additional purification steps allowed to separate two group of diastereoisomers, LJ2a and LJ2b, harboring stereoselectivity in P2. We then decided to synthetize LJ3 and adopted a similar purification approach to separate the P2-stereoselective diastereoisomers LJ3a and LJ3b.

Then, we have determined the kinetic parameters k_3_/K_i_, k_3_, and K_i_, for LJ2a, LJ2b, LJ3a, and LJ3b on human Casp2 and Casp3 (Table 1). LJ2a is a highly potent Casp2 inactivator showing a subnanomolar K_i_ for Casp2 and a k_3_/K_i_ ratio of ~ 5 500 000 M^−1^s^−1^(i.e. 400% improvement as compared to LJ2). LJ2a also inactivates Casp3 but ~50 times less than Casp2 (LJ2a^Casp3^k_3_/K_i_ = 113 425 M^−1^s^−1^). Spatial orientation of the 6-methyl-1,2,3,4-tetrahydroisoquinoline-1-carbonyl residue (P2 position) appeared crucial for Casp2 inactivation as the k_3_/K_i_ ratio of LJ2b against Casp2 is 765 times lower than the one of LJ2a.

**TABLE 1:** Caspase inhibition activity of P2-stereospecific LJ2a, LJ2b, LJ3a, LJ3b compounds against Casp-2 vs Casp-3.

| Compounds | Casp2 |  |  | Casp3 |  |  |
| --- | --- | --- | --- | --- | --- | --- |
| | $k_3$<br>($\text{ms}^{-1}$) | $K_i$<br>(nM) | $k_3/K_i$<br>($\text{M}^{-1}.\text{s}^{-1}$) | $k_3$<br>($\text{ms}^{-1}$) | $K_i$<br>(nM) | $k_3/K_i$<br>($\text{M}^{-1}.\text{s}^{-1}$) |
| LJ2a | $5 \pm 5 \times 10^{-4}$ | $0.9 \pm 0,1$ | 5 499 419 | $1.3 \pm 4 \times 10^{-5}$ | $11,5 \pm 0.9$ | 113 425 |
| LJ2b | $4 \pm 3 \times 10^{-4}$ | $524 \pm 10$ | 7 177 | $8 \pm 2 \times 10^{-3}$ | $17\,498 \pm 104$ | 457 |
| LJ3a | $5 \pm 7 \times 10^{-4}$ | $2.8 \pm 0.8$ | 1 708 996 | $0.7 \pm 9 \times 10^{-5}$ | $399,8 \pm 93,3$ | 1 807 |
| LJ3b | $7 \pm 10^{-3}$ | $8\,539 \pm 300$ | 823 | $0,8 \pm 6 \times 10^{-5}$ | $84\,205 \pm 13\,000$ | 9 |

LJ3a is also an excellent inactivator of Casp2 showing a 2.8nM K_i_ and k_3_/K_i_ ratio of ~ 1 709 000 M^−1^s^−1^ vis-a-vis Casp2. Comparatively, LJ3a has a low inactivation rate on Casp3 (946 times less than for Casp2) (LJ3a^Casp3^ k_3_/K_i_ = 1 807 M^−1^s^−1^). Spatial orientation of the 3-neopentyl-Proline residue (P2 position) appeared crucial for Casp2 inactivation as the k_3_/K_i_ ratio of LJ3b against Casp2 was more than 2000 times lower than the one of LJ3a.

Additional enzyme kinetics experiments with human recombinant Caspase-1, Caspase-6, Cathepsin-B, Cathepsin-L, Cathepsin-D, Thrombin, Plasmin, Trypsin, Kallikrein-1, −6, and −8 indicates that LJ2a and LJ3a have very limited or no effect on these enzymes at least when used at concentrations up to 10μM. The only observed effect was a weak Casp-6 inhibition by LJ2a (LJ2a^Casp6^ k_3_/K_i_ = 14.2 M^−1^s^−1^) but not by LJ3a. Hence, LJ2a is the most potent Caspase-2 inhibitor ever reported, and it is a mild inhibitor of Caspase-3. LJ3a is both a highly potent and genuinely selective Casp2 inhibitor.

### Structural Characterization of LJ3a and LJ3b

Consequently, we decided to further characterize the LJ3a and LJ3b compounds by NMR and molecular modeling. ^1^H-^15^N SOFAST-HMQC experiments in natural abundance allowed to observe each peak corresponding to one amino acid at positions P5 (Val 5), P4 (Asp 4), P3 (Val 3) and P1 (Asp-OPh). No signal can be observed for the position P2 (3-neopentyl proline) since the nitrogen of the proline does not carry a proton. The ^1^H-^15^N SOFAST-HMQC experiments in natural abundance performed on the LJ3a and LJ3b samples showed a doubling of some resonances demonstrating that they did not contain a single isomer but a mixture of 2 different isomers (Fig. 6a,b). These results were confirmed by 1H NMR experiments performed on these samples (Suppl. Fig. S1A, S2A). The resonances corresponding to the amide group N6H36 in position P1 (ASP 1) are split in each ^1^H-^15^N SOFAST-HMQC experiment in natural abundance performed on the samples LJ3a and LJ3b. The resonances of the amide group N2H7 corresponding to position P5 (VAL 5) and amide group N3H16 at position P4 (ASP 4) are unique in the 2 samples confirming the absolute configuration S is unchanged for the 2 α-carbons C11 and C16 at position P5 and P4 respectively (Fig. 6a,b). The two peaks N6H36 corresponding to the two isomers at position P1 appeared to be in a 50/50 ratio when measuring the volume of the correlation peaks for each sample. This indicated a racemization of the α-carbon C30 at the position P1. In both samples, LJ3a and LJ3b, the resonances of the amide group N4H21 at position P3 (Val 3) showed the presence of 2 signals. However, the absolute configuration S of this carbon should remain unchanged due to the solid phase synthesis mode used and it cannot be ruled out that the electronic environment of the H21 is changed due to the racemization of the position P1 (ASP 1), resulting in the doubling of the signals of the amide group. The N5 of proline 2 in position P2 does not carry a proton and is therefore not observable on the ^1^H-^15^N SOFAST-HMQC experiments. Nevertheless, the Hα (H32) of the 3-neopentyl proline in position P2 shows two distinct resonances showing that it undergoes an effect due to the presence of the two isomers in position P1. Thus, the α-carbon at the position P1 is racemic and position P2 is optically pure. Then, the full assignment of the isomers and exact structures of LJ3a (Fig. 6c-e) and LJ3b (Fig. 6f-h) were established using ^1^H NMR experiments and comparison of spectra with theoretical models (Suppl. Fig S1, S2, Suppl Table 1, 2).

### LJ2a and LJ3a protect primary neurons against β-amyloid-induced synapse loss

Caspase-2 is critical for the effects of Aβ on dendritic spines in cultured neurons and has a critical role in mediating the synaptic changes and memory alteration induced by Aβ in human amyloid precursor protein transgenic mice, suggesting that Caspase-2 is a potential target for Alzheimer’s disease therapy [17]. We cultured hippocampal neurons and exposed them to low (subapoptotic) concentrations of oligomeric Aβ_1-42_ (nM range) with and without Caspase-2 inhibitors and examined the effects on dendritic spine density (Fig.7). This resulted in a 50% decrease in spine density after 6h of exposure. These effects were not seen in neurons exposed to monomeric Aβ_1-42_. When submicromolar concentration of LJ2a were added, there was a full blockade of the effect of oligomeric Aβ on spine density (Fig 7b). LJ3a pretreatment at 0.1 μM, 1 μM, and 10 μM prevented dendritic spine loss induced by oligomeric Aβ at 59.63%, 89.29%, and 97.5% respectively (Fig 7c).

**Figure 7:**
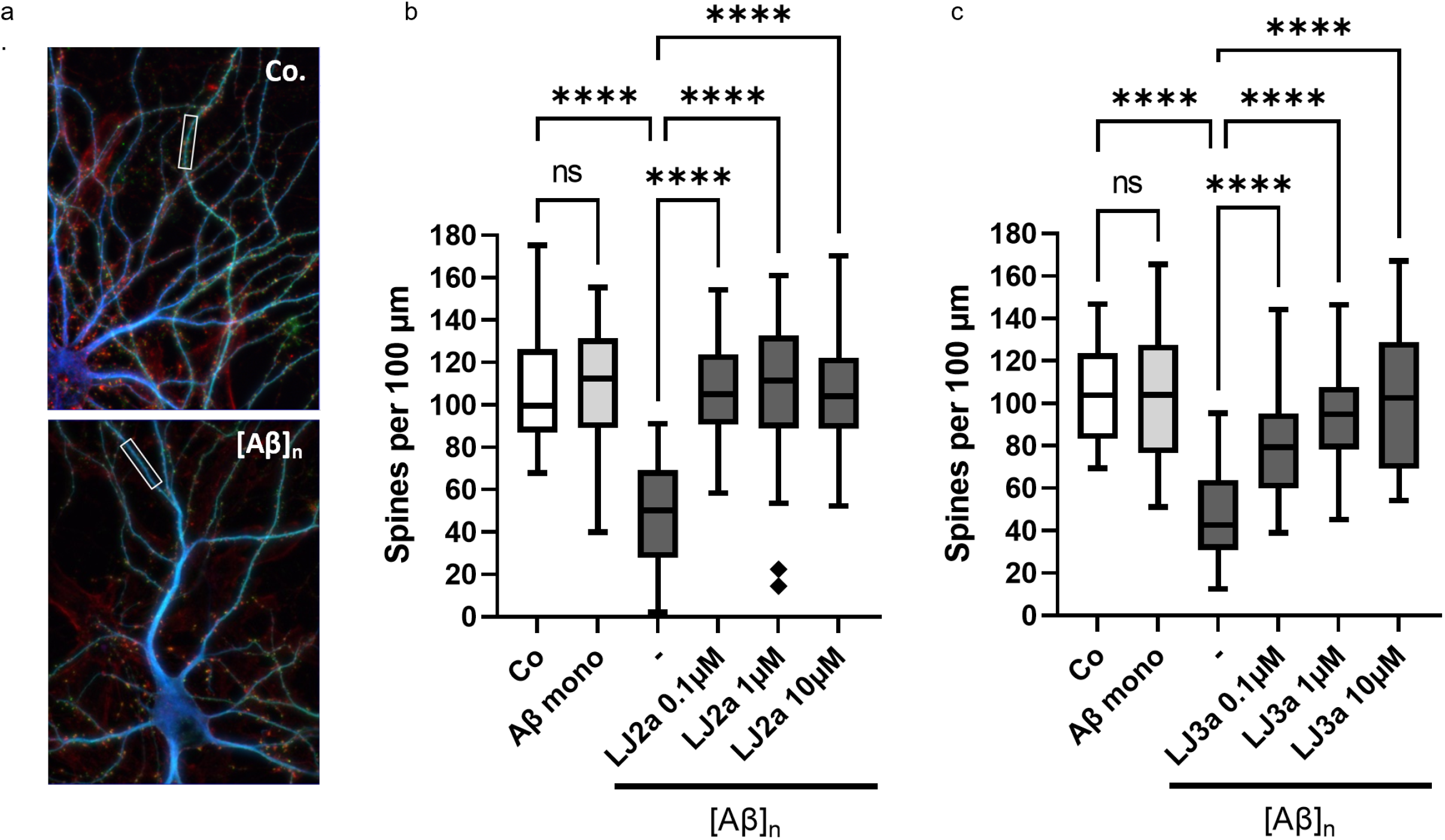
LJ2a and LJ3a prevent dendritic spine loss induced by Aβ oligomers in hippocampal primary cultures. Swiss mice hippocampal neurons (E18) were cultured for 3 weeks in microfluidic chambers (20 000 neurons / chamber). Then neurons were pre-treated, or not, for 1h with the indicated concentration of LJ2a or LJ3a, and treated for 6h with 100 nM of monomeric Aβ (Aβ mono) or 100 nM of Aβ_1-42_ oligomers ([Aβ]_n_) then fixed and permeabilized for (immuno)staining with anti- MAP2, Phalloidin, anti-actin F, and anti-Bassoon. Microfluidic chambers were analyzed by fluorescence microscopy by counting phalloidin clusters affixed to MAP2 and Bassoon on hippocampal dendrites. **a.**Representative micrographs of triple stained dendrites after 6h in the presence (lower panel, [Aβ]_n_) or absence (upper panel, Co.) of Aβ oligomers. **b-c.** Quantification of dendritic spines in hippocampal neurons treated with LJ2a (b) or LJ3a (c) as indicated. Both inhibitors show synaptoprotective effects at submicromolar concentration. Histograms represent means (+/-SD) of 3 independent experiments (****p value < 0.0001).

### LJ2a and LJ3a abolish Casp2-dependent SREBP2 activation

Previously, it was found that Casp2 activates SREBP1 and 2, the master regulators of lipogenesis and cholesterol biosynthesis, through the cleavage-mediated activation of S1P [12]. Casp2 inhibition reduces hepatic steatosis and steatohepatitis, inflammation, and liver damage, as well as it prevents cholesterol accumulation and hyperlipidemia, suggesting that Casp2 is an interesting target for the prevention or treatment of NASH [12]. To evaluate Casp2 inhibitor effects on SREBP activation, we added the inhibitors to HEK293 cells transfected with Casp2, S1P, and SREBP2 (Fig. 8a, b). Under these conditions, Casp2 cleaves and activates S1P which then cleaves and activates SREBP2 resulting in nuclear translocation of the SREBP2 transcriptional activation domain [12] (Fig. 8b, Lane 2). Strikingly, addition of LJ2a or LJ3a completely abolished Casp2- and S1P-dependent SREBP2 activation (Fig. 8b, Lane 4, 6) while TRP601 led to partial inhibition (Lane 8).

**Figure 8:**
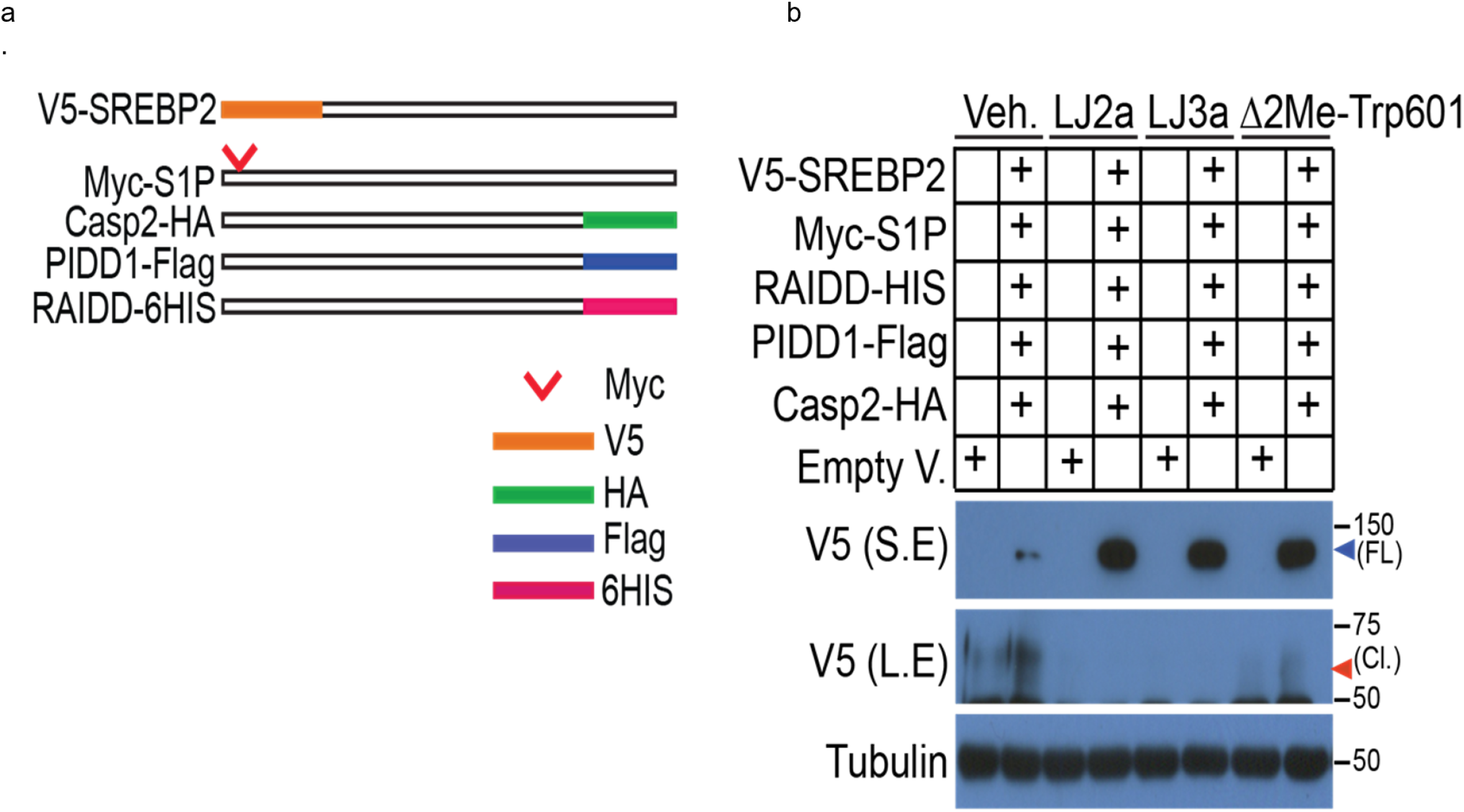
Caspase-2 inhibitors prevent sterol regulatory element-binding protein 2 (SREBP2) activation via the inhibition of site 1 protease (S1P) cleavage. **a.** Schematic representation of cDNAs encoding HA-tagged Casp2, Myc-tagged S1P at the carboxy terminus (S1P-Myc), V5-tagged SREBP2, PIDD1-Flag, RAIDD-6HIS. Intracellular S1P was detected by an S1P antibody that recognizes the first 50 AA and membrane and secreted S1P polypeptides were detected by an antibody that recognizes AA200 to AA300. **b.** HEK 293 cells were transfected with indicated plasmids in order to overexpress SREBP2, S1P, RAIDD, PIDD, and Casp-2. After 5 hrs, cells were incubated in DMEM/F12 medium supplemented with vehicle, LJ2a, LJ3a, or Δ2ME-TRP601 (10 μM) for 16h. Then whole cell extracts where subjected to WB to detect (cleaved) SREBP2 using an antibody directed against the V5 epitope.

## Discussion

Caspases are implicated in the pathogenesis of numerous diseases. Hundreds of caspase-specific inhibitors have been designed, few have progressed to clinical trials, and to date no one has reached market authorization. This is due, at least in part, to the use of non-selective approaches to caspase inhibition which inhibit the whole family rather than the key family member responsible for pathology. Designing highly selective and druggable reagents to distinguish among closely related enzymes of the caspase family remains a major challenge. This difficulty is exacerbated for the design of active-site-directed inhibitor of Casp2 due to high similarity between Casp3 and Casp2 active sites.

We have designed and evaluated a series of peptidomimetics inspired from the pentapeptide Val-Asp-Val-Ala-Asp (VDVAD) and included N-terminal quinaldoyl group, potent irreversible C-terminal warhead (proven to be safe in human), and chosen non-natural bulky structures in P2 to generate new compounds harboring higher biodisponibility, improved efficiency, and enhanced selectivity toward Caspase-2.^31-33^ Our results define a series of highly potent Casp2 inhibitors having the general structure: Quinaldoyl-Val^P5^-Asp^P4^-Val^P3^-X^P2^-Asp^P1^-fluorophenoxy-methyl-ketone (*Q*-VDVXD-*OPh*) with X^P2^ being either 6-methyl-tetrahydro-isoquinoline (LJ2 and LJ2a) or a 3-(S)-neopentyl proline (LJ3a). LJ2 has high inactivation rate on Casp2 (k_3_/K_i_ ~ 1 300 000 M^−1^s^−1^) and shows selectivity (~25 times higher as compared to Casp3). In cell lines, LJ2 dose-dependently inhibits cell death induced by microtubule destabilization or hydroxamic acid-based deacetylase inhibition, two Casp2-dependent cell deaths paradigms. LJ2a (further purified from LJ2), is the most active compound. It has a subnanomolar K_i_ for Casp2, a very high inactivation rate on Casp2 (k_3_/K_i_ ~ 5 500 000 M^−1^s^−1^) and show good selectivity (~50 times higher as compared to Casp3). The most selective compound, LJ3a, has a high inactivation rate on Casp2 (k_3_/K_i_ ~ 1 700 000 M^−1^s^−1^) and high selectivity (~1000 times higher as compared to Casp3). Enzyme kinetics show that LJ3b isomer has very low activity on Casp2 (k_3_/K_i_ ~ 800 M^−1^s^−1^). Structural analysis of LJ3a and LJ3b shows that precise spatial configuration R/S of the α-carbon in P2 determines inhibitor efficacy. Indeed, NMR spectra analysis and molecular modeling show that the position P2 (3-neopentyl proline) is optically pure in LJ3a whereas the α-carbon in P1 is racemic. LJ3a contains the two isomers (P1_R_, P2_R_) and (P1_S_, P2_R_) while sample LJ3b contains the two isomers (P1_R_, P2_S_) and (P1_S_, P2_S_). Hence, the unexpected conclusion is that the S-configuration of the α-carbon in P2 (LJ3b) is inactive against Casp2, whereas the R-configuration of the α-carbon in P2 (LJ3a) is active.

Casp2 is implicated in several diseases including optic nerve injuries, neonatal brain damage, age-related neurodegeneration, and metabolic diseases. To investigate the effect of LJ2a and LJ3a in pathological conditions, we selected two diseases, AD and NASH, because they are both devastating diseases with a huge world-wide societal impact and so far no drug have shown disease-modifying efficacy against either of them [42,43].

Casp2 is a potential therapeutic target in AD [17, 18]. Indeed, experiments with primary hippocampal neurons and Casp2-deficient mice implicate Casp2 as key driver of synaptic dysfunction and cognitive decline in AD [17]. Casp2 is present both in neuron cell bodies and dendritic spines and acts as a mediator of β-amyloid protein (Aβ) synaptotoxicity. Consequently, one can expect that selective Casp2 inhibitors would inhibit Aβ-induced synapse loss. The present study shows that, in primary hippocampal neurons, submicromolar concentrations of LJ2a and LJ3a block synapse loss induced by Aβ42 oligomers. The preclinical development of the (less selective) parent compound, TRP601, was previously reported [31]. It was found to be non-toxic in regulatory rodent and non-rodent studies, to inhibit of Casp2 in the brain, and to confer neuroprotection after ip and iv administration [31]. One can expect that LJ2a and LJ3a may have such favorable safety and PK properties. Further studies in human cells and animal models of AD are ongoing to investigate the potential of LJ2a and LJ3a for the treatment of AD.

The first in class dipeptide derivative irreversible pan-caspase inhibitor, Emricasan, has been thoroughly investigated in several clinical studies for a variety of liver diseases. This drug candidate, recently failed to demonstrate efficacy in large clinical phase 3 studies for the treatment of NASH [44, 45]. Those studies however, provided important information that contradicted broadly held opinion, showing that chronic administration of an irreversible broad-spectrum Caspase inhibitor is not toxic nor carcinogenic in humans (Clinical Trials.gov #NCT02686762 and #NCT02960204) [44, 45]. This paves the way for the development of more selective drugs directed against individual Caspases. Other recent studies in cellular and animal models, have highlighted the importance of Casp2, in NASH progression [12]. Indeed, Casp2 inhibition may lead to reduced lipoapoptosis and steatohepatitis, and block the production of fibrogenic Hedgehog ligands, which aggravate NASH progression [11, 46]. More recently, it was suggested that Casp2 activation is a critical mediator of the transition from benign non-alcoholic fatty liver disease (NAFLD) to NASH [12]. Casp2 activation was associated with dysregulated SREBP1/2 activation, which is accompanied by lipid and cholesterol accumulation within the liver [14]. Altogether these findings imply that Casp2 inhibition could be a valid therapeutic approach to stop the pathogenic progression that leads to NASH. Considering the failure of Emricasan clinical results, we suggest that successful prevention or treatment of NASH would require the use of a Casp2-specific inhibitor. Our data show that LJ2a and LJ3a inhibit the Casp2 mediated processing of SREBP2, in a cellular model. This suggests that LJ3a and LJ2a should be further investigated and optimized for *in vivo* activity to determine if selective Casp2 inhibitors could offer an effective approach to the prevention or treatment of fatty liver diseases.

## Acknowledgements

We thank Dr Michel Maillard (CHDI) for advice and kindly providing reversible Caspase-2 inhibitors (q33, h33, k33 and c33). Elodie Bosc was funded by MESNR. We thank Ségolène Prétat and Hugot Cochet for technical assistance. This work was funded by ANR “Neuroscreen: 2011-RPIB-008-001” 2012-16 (to BB), SATT IDF Maturation grant (to EJ), iXLife-iXCore-iXBlue Fondation pour la Recherche (to EJ), and NIH grant DK120714 (to MK).

## Declaration of interests

E.J. is inventor of a patent titled “Novel derivatives and their use as selective inhibitors of Caspase-2” (WO/2017/162674). E.J. and E.B. are inventors of a patent titled “Novel compounds and their use as selective inhibitors of Caspase-2” WO/2019/068538. MK hold a patent on mouse models for the treatment of NASH, is a founder of Elgia Pharmaceuticals and had received research support from Jansen and Gossamer Bio.

