## Supplemental Methods for "Genuine Selective Caspase-2 Inhibition with new Irreversible Small Peptidomimetics"

### Supplementary Data

#### Supplementary Figures

##### Figure S1

(A) Superposition of the NH / H$\alpha$ regions of the NOESY (cyan) and TOCSY (magenta) spectra recorded on the LJ3a sample. The superimposition of the two spectra makes it possible to identify each amino acid by the TOCSY experiment and the sequential succession of these residues by the NOESY experiment. Sequential assignment is indicated by horizontal and vertical lines. The resonances are doubled for the positions P3 (Val 3) and P1 (Asp 1), then two sets of data can be identified, labeled in thin and in bold lettering and corresponding to two different isomers. (B) Table of chemical shifts $^{15}$N and $^{1}$H of the skeleton and side chains of the different residues of LJ3a. It is found that the chemical shift values are similar for the protons of the side chains and different for the $^{15}$N, HN and H$\alpha$.

##### Figure S2

(A) Superposition of the NH / H$\alpha$ regions of the NOESY (cyan) and TOCSY (magenta) spectra recorded on the LJ3b sample. The superimposition of the two spectra makes it possible to identify each amino acid by the TOCSY experiment and the sequential succession of these residues by the NOESY experiment. Sequential assignment is indicated by horizontal and vertical lines. The resonances are doubled for the positions P3 (Val 3) and P1 (Asp 1), then two sets of data can be identified, labeled in thin and in bold lettering and corresponding to two different isomers. (B) Table of chemical shifts $^{15}$N and $^{1}$H of the skeleton and side chains of the different residues of LJ3b. It is found that the chemical shift values are similar for the protons of the side chains and different for the $^{15}$N, HN and H$\alpha$.

#### Supplementary Tables

##### Suppl. Table 1

Comparison of the distances measured between protons on each 3D theoretical model of the 1R2R, 1S2R, 1R2S and 1S2S isomers and the volumes and distances derived from the

NOESY experiment at 400 ms between these same protons. This comparison made it possible to identify the 2R or 2S isomer present in each sample and to conclude that LJ3a contains isomer 1(R,S)2R and that LJ3b contains isomer 1(R,S)2S. (*) indicates that two signals are superimposed.

**Suppl. Table 2**

Comparison of the distances measured between protons on each 3D theoretical model of the 1R2R, 1S2R, 1R2S and 1S2S isomers and the volumes and distances derived from the NOESY experiment at 100 ms between these same protons. This comparison made it possible to identify the 2R or 2S isomer present in each sample and to conclude that LJ3a contains isomer 1(R,S)2R and that LJ3b contains isomer 1(R,S)2S. (∅) indicates that no signal is detectable.

**Suppl. Table 1**

Comparison of the Measured distances on the model structures with the measured volumes and derived distances from the 400 ms NOESY spectra.

| Measured distance on the model structures (Å) | | | | | | Measured volumes and derived distances on 400 ms NOESY spectra | | | | Result |
|---|---|---|---|---|---|---|---|---|---|---|
| Protons | | 1R2R | 1S2R | 1R2S | 1S2S | LJ3a1 | LJ3a2 | LJ3b1 | LJ3b2 | |
| H21 | H30 | 4.39 | 4.28 | 1.80 | 2.62 | 550018<br>4.43 | * | 820696<br>3.90 | 693644<br>4.00 | LJ3b=1(R,S)2S |
| H21 | H31 | 5.21 | 5.14 | 3.27 | 3.70 | 294592<br>4.92 | 354974<br>4.77 | 1050000<br>3.74 | 976086<br>3.74 | LJ3b=1(R,S)2S |
| H22 | H35 | 5.51 | 5.61 | >6.0 | >6.0 | 481831<br>4.04 | 481831<br>4.04 | 320176<br>4.06 | 320176<br>4.06 | LJ3a=1(R,S)2R |
| H30 | H46 | 4.89 | 4.87 | 4.00 | 4.08 | 110876<br>5.16 | 110876<br>5.16 | 96120<br>4.96 | 96120<br>4.96 | LJ3b=1(R,S)2S |
| H31 | H46 | 4.48 | 4.39 | 2.58 | 2.63 | 214238<br>4.63 | 214238<br>4.63 | 800000<br>3.49 | 800000<br>3.49 | LJ3b=1(R,S)2S |
| H32 | H37 | 4.21 | 4.07 | 4.39 | 4.23 | 520824<br>4.48 | 656653<br>4.31 | 346066<br>4.50 | 470582<br>4.28 | LJ3a=1(R,S)2R |
| H33 | H36 | 3.76 | 3.92 | 2.52 | 3.16 | 554349<br>4.43 | 801356<br>4.17 | 801016<br>3.91 | 1010000<br>3.76 | LJ3b=1(R,S)2S |
| H34 | H37 | >6.0 | 5.69 | 5.55 | 4.30 | 239313<br>5.00 | 231267<br>5.13 | 113630<br>5.00 | 100462<br>5.50 | LJ3a=1(R,S)2S |
| H37 | H46 | 4.68 | 2.36 | >6.0 | 5.96 | 89113<br>6.00 | 94410<br>5.96 | 65388<br>5.94 | 67752<br>5.90 | LJ3a=1(R,S)2R |
| H37 | H47 | 4.79 | 2.67 | 5.91 | 4.94 | 311039<br>4.88 | 294443<br>4.93 | 131594<br>5.29 | 190282<br>4.97 | LJ3a=1(R,S)2R |

* indicates two superimposed signals

**Suppl Table 2**

Comparison of the Measured distances on the model structures with the measured volumes and derived distances from the 100 ms NOESY spectra

| Measured distance on the model structures (Å) | | | | | | Measured volumes and derived distances from the 100 ms NOESY spectra | | | | Result |
|---|---|---|---|---|---|---|---|---|---|---|
| Protons | | 1R2R | 1S2R | 1R2S | 1S2S | LJ3a1 | LJ3a2 | LJ3b1 | LJ3b2 | |
| H21 | H30 | 4.39 | 4.28 | 1.80 | 2.62 | 179675<br>4.50 | ∅ | 107604<br>3.92 | 103660<br>3.95 | LJ3b=1(R,S)2S |
| H21 | H31 | 5.21 | 5.14 | 3.27 | 3.70 | 81438<br>3.85 | 58925<br>4.07 | 135778<br>3.77 | 129023<br>3.80 | LJ3b=1(R,S)2S |
| H22 | H35 | 5.51 | 5.61 | >6.0 | >6.0 | 37858<br>3.90 | 37858<br>3.90 | 22293<br>3.77 | 22293<br>3.77 | LJ3a=1(R,S)2R |
| H30 | H46 | 4.89 | 4.87 | 4.00 | 4.08 | ∅ | ∅ | 17798<br>4.71 | 17798<br>4.71 | LJ3b=1(R,S)2S |
| H31 | H46 | 4.48 | 4.39 | 2.58 | 2.63 | 30726<br>4.04 | 30726<br>4.04 | 116236<br>3.45 | 116236<br>3.45 | LJ3b=1(R,S)2S |
| H32 | H37 | 4.21 | 4.07 | 4.39 | 4.23 | 73513<br>3.96 | 88556<br>3.80 | 35391<br>4.72 | 22142<br>5.10 | LJ3a=1(R,S)2R |
| H33 | H36 | 3.76 | 3.92 | 2.52 | 3.16 | 83350<br>3.84 | 103640<br>3.70 | 91508<br>4.03 | 129888<br>3.8 | LJ3b=1(R,S)2S |
| H34 | H37 | >6.0 | 5.69 | 5.55 | 4.30 | 27434<br>4.39 | 34243<br>4.45 | ∅ | 20158<br>5.18 | LJ3a=1(R,S)2S |
| H37 | H46 | 4.68 | 2.36 | >6.0 | 5.96 | ∅ | ∅ | ∅ | ∅ | |
| H37 | H47 | 4.79 | 2.67 | 5.91 | 4.94 | 37717<br>4.38 | 42793<br>4.29 | ∅ | ∅ | LJ3a=1(R,S)2R |

∅ no detectable signal

#### Supplementary methods

##### Full assignment of the isomers in LJ3a and LJ3b using $^{1}$H NMR

To determine which sample contained the 2R and which one contains the 2S, we performed the full assignment of the isomers using $^{1}$H NMR experiments (1D $^{1}$H, 2D TOCSY, 2D NOESY, 2D COSY) (Suppl. Fig. S1, S2). We observed that chemical shifts of the protons involved in the racemization were different on the spectra of the two samples LJ3a and LJ3b. For example, H36/H4* was observable for one given isomer as a single correlation peak, while for the other isomer of the same sample, two correlation peaks, H36/H41 and H36/H42, were observed. In contrast, the chemical shifts of the side chains protons of the different isomers were similar in the two batches LJ3a and LJ3b and were therefore not sensitive to changes in the electronic environment of the protons attached to the carbons with the absolute configuration R or S (Figure S1B, S2B). The most important chemical shifts modifications concerned the nuclei N (N6), NH (H36) and H$\alpha$ (H37) of the racemic residue D1 in the two sample LJ3a and LJ3b confirming the racemization of the position P1. To identify which of the isomers (1R, 2R), (1S, 2R) on the one side and (1R, 2S), (1S, 2S) on the other side, could correspond to the recorded spectra, we built a theoretical model of each of these isomers. Then we measured on these theoretical models the characteristic inter-proton distances that discriminate the two isomers in position P2 (Suppl. Table 1, Suppl. Table 2). In a second step, we measured on the NOESY spectra, recorded at two different mixing times to account for spin diffusion artifacts, the volumes of the signals corresponding to the distances between protons measured previously on the theoretical models. The experimental distances were then determined from these volumes. Finally, knowing that the volume of the NOE between two nuclei across space depends inversely on the distance separating these two nuclei, we compared the volume of the peaks measured on the NOESY spectra, and the derived distances, to the distance determined on the theoretical models to determine which had the best match between experimental volume and theoretical distance. Ten distances were measured on the theoretical models of the two isomers (1R, 2R), (1S, 2R) and (1R, 2S), (1S, 2S) and compared to the distances deduced from the NOE volumes. Of these 10 distances, 9 show that sample LJ3a contains the two isomers (1R, 2R) and (1S, 2R) and that sample LJ3b contains the two isomers (1R, 2S) and (1S, 2S).

**Non-caspase enzyme assays**

The human pro-kallikrein-1 (pro-hK1) was activated in a buffer containing: Tris-HCl 50 mM, CaCl2 10 mM, NaCl 150 mM, Brij-35 0,05% pH 7,5. Human pro-kallikrein-6 and -8 (pro-hK6 and 8 were activated using Lysyl-endopeptidase (Wako-BioProducts®) at a 1/500 weight ratio, in a buffer containing 50 mM Tris and 0,05% Brij-35 (pH 8). Kinetics using non-caspase enzymes for selectivity studies were done according to the following conditions.

| **Enzymes** | **Substrates** | **Buffers** |
|---|---|---|
| Cathepsine B (0.2 nM) | z-RR-AMC 20 µM<br>($K_M$ = 169,8 µM) | Acétate de sodium 0,1 M ; EDTA 1mM ;<br>DTT 2mM ; Brij-35 0,01 % ; pH 5,5 |
| Cathepsine L (1.2 nM) | RLR-AMC 25 µM<br>($K_M$ = 13,8 µM) | Acétate de sodium 0,1 M ; EDTA 1mM ;<br>DTT 2mM ; Brij-35 0,01 % ; pH 5,5 |
| Cathepsine D (0.1 nM) | Mca-Gly-Lys-Pro-Ile-Leu-Phe-Phe-Arg-Leu-Lys(Dnp)-D-Arg-$NH_2$ 10 µM | Citrate de Sodium 0,1 M ; EDTA 2 mM ;<br>Brij-35 0,01% ; pH 4 |
| hK1 (4 nM) | Boc-VPR-AMC 100µM | Tris 50mM; Citrate 1M; Brij-35 0.05%; pH7 |
| hK6 (2 nM) | Boc-QAR-AMC 100 µM | Tris 50mM; Citrate 1M; Brij-35 0.05%; pH7 |
| hK8 (1 nM) | Boc-VPR-AMC 100µM | Tris 50mM; Citrate 1M; Brij-35 0.05%; pH7 |
| Thrombine (10 mU/mL) | Boc-VPR-AMC 100µM | Tris 50mM; Citrate 1M; Brij-35 0.05%; pH7 |
| Plasmine (4 nM) | Boc-QAR-AMC 100µM | Tris 50mM; Citrate 1M; Brij-35 0.05%; pH7 |
| Trypsine (0.1 nM) | Boc-QAR-AMC 100µM | Tris 50mM; Citrate 1M; Brij-35 0.05%; pH7 |
