## Supplemental Figure S2 for "Genuine Selective Caspase-2 Inhibition with new Irreversible Small Peptidomimetics"

A

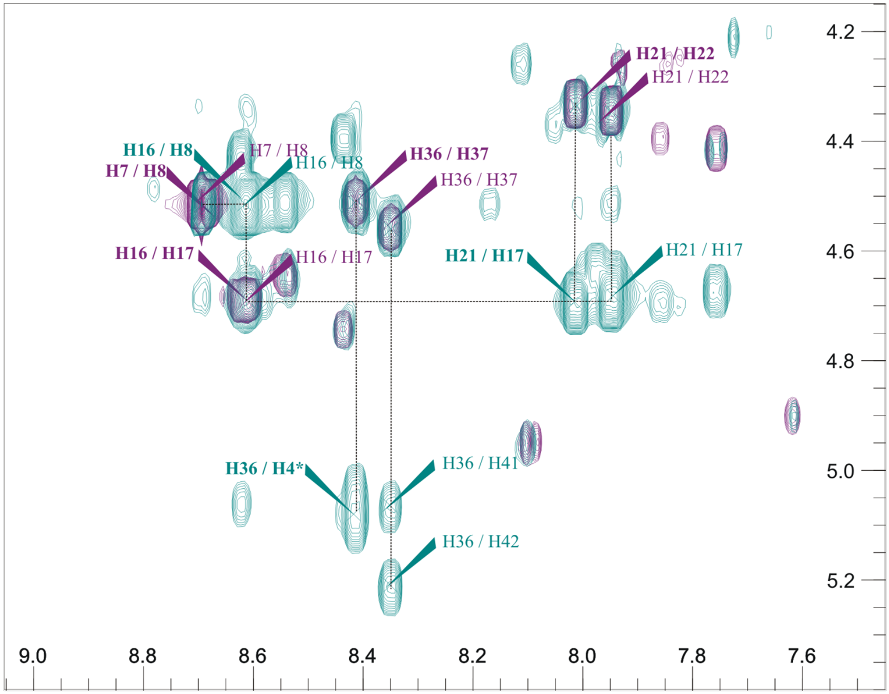


B

| Residue | N | NH | αH | βH | Others |
|---|---|---|---|---|---|
| **D1** | 114.2 | 8.41 | 4.51 | 2.49, 2.84 | 5.08 |
| | 113.71 | 8.35 | 4.56 | 2.53, 2.72 | 5.06, 5.21 |
| **X2** | | | 3.69 | 2.04 | γ$CH_2$: 1.49, 2.12; δ$CH_2$: 3.50, 3.67; ε: 1.21, 1.49 |
| | | | 3.74 | 2.06 | γ$CH_2$: 1.49, 2.12; δ$CH_2$: 3.50, 3.67; ε: 1.21, 1.49 |
| **V3** | 115.64 | 8.02 | 4.33 | 1.97 | Hγ1# 0.79 ; Hγ2# 0.79 |
| | 115.42 | 7.96 | 4.35 | 1.97 | Hγ1# 0.79 ; Hγ2# 0.79 |
| **D4** | 120.28 | 8.61 | 4.69 | 2.55, 2.69 | |
| **V5** | 106.92 | 8.69 | 4.51 | 2.15 | Hγ1# 0.90 ; Hγ2# 0.90 |

**Supplementary Figure S2. Bosc et al., 2021**
