## Supplemental Figure S1 for "Genuine Selective Caspase-2 Inhibition with new Irreversible Small Peptidomimetics"

A

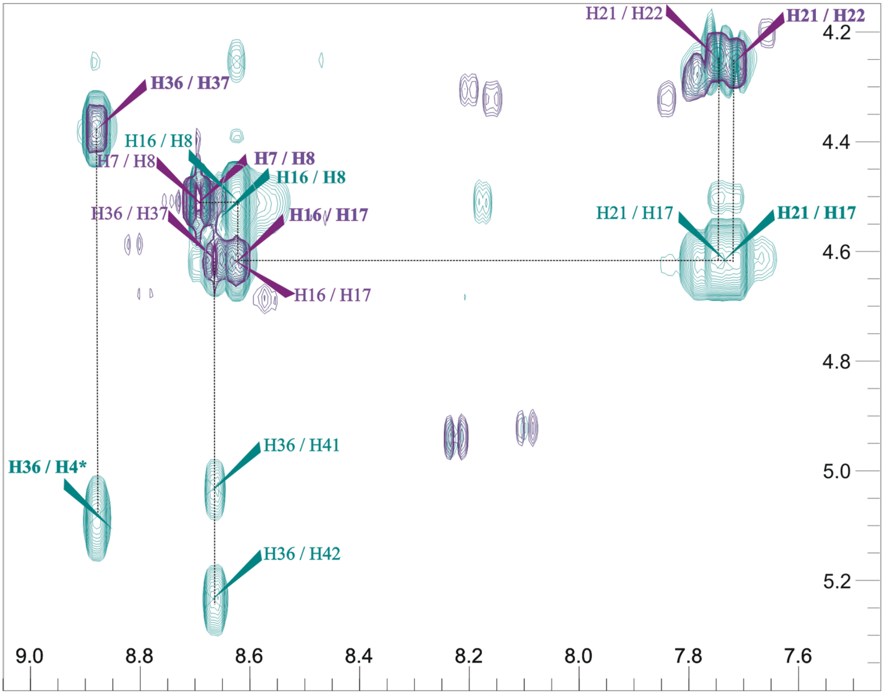


B

| **Residue** | **N** | **NH** | **αH** | **βH** | **Others** |
|---|---|---|---|---|---|
| **D1** | 117.07 | 8.88 | 4.38 | 2.49, 2.84 | 5.08 |
| | 118.01 | 8.66 | 4.61 | 2.53, 2.72 | 5.03, 5.23 |
| **X2** | | | 3.67 | 2.06 | γ$CH_2$: 1.48, 2.20; δ$CH_2$: 3.47, 3.84; ε: 1.21, 1.49 |
| | | | 3.70 | 2.04 | γ$CH_2$: 1.50, 2.19; δ$CH_2$: 3.47, 3.84, ε: 1.21, 1.49 |
| **V3** | 116.15 | 7.72 | 4.25 | 1.81 | Hγ1# 0.76 ; Hγ2# 0.91 |
| | 116.24 | 7.75 | 4.25 | 1.82 | Hγ1# 0.76 ; Hγ2# 0.76 |
| **D4** | 120.64 | 8.62 | 4.72 | 2.50, 2.65 | |
| **V5** | 107.15 | 8.69 | 4.51 | 2.13 | Hγ1# 0.93 ; Hγ2# 0.93 |

**Supplementary Figure S1. Bosc et al., 2021**
